# Clinical Knowledge Graph Integrates Proteomics Data into Clinical Decision-Making

**DOI:** 10.1101/2020.05.09.084897

**Authors:** Alberto Santos, Ana R. Colaço, Annelaura B. Nielsen, Lili Niu, Philipp E. Geyer, Fabian Coscia, Nicolai J Wewer Albrechtsen, Filip Mundt, Lars Juhl Jensen, Matthias Mann

**Affiliations:** NNF Center for Protein Research, Faculty of Health Sciences, University of Copenhagen, 2200 Copenhagen, Denmark; Department of Proteomics and Signal Transduction, Max Planck Institute of Biochemistry, 82152 Martinsried, Germany; OmicEra Diagnostics GmbH, Planegg, Germany; Department of Biomedical Sciences, Faculty of Health and Medical Sciences, University of Copenhagen, Copenhagen, Denmark; Department for Clinical Biochemistry, Rigshospitalet, University of Copenhagen, Copenhagen, Denmark; Li-Ka Shing Big Data Institute, University of Oxford, UK

**Author notes:** Correspondence: Alberto Santos or Matthias Mann.

**Keywords:** network biology, multi-omics, data integration, systems biology, mass spectrometry, python, jupyter notebooks, machine learning, liver disease, oncology

## Abstract

The promise of precision medicine is to deliver personalized treatment based on the unique physiology of each patient. This concept was fueled by the genomic revolution, but it is now evident that integrating other types of omics data, like proteomics, into the clinical decision-making process will be essential to accomplish precision medicine goals. However, quantity and diversity of biomedical data, and the spread of clinically relevant knowledge across myriad biomedical databases and publications makes this exceptionally difficult. To address this, we developed the Clinical Knowledge Graph (CKG), an open source platform currently comprised of more than 16 million nodes and 220 million relationships to represent relevant experimental data, public databases and the literature. The CKG also incorporates the latest statistical and machine learning algorithms, drastically accelerating analysis and interpretation of typical proteomics workflows. We use several biomarker studies to illustrate how the CKG may support, enrich and accelerate clinical decision-making.

**Graphical Abstract:** 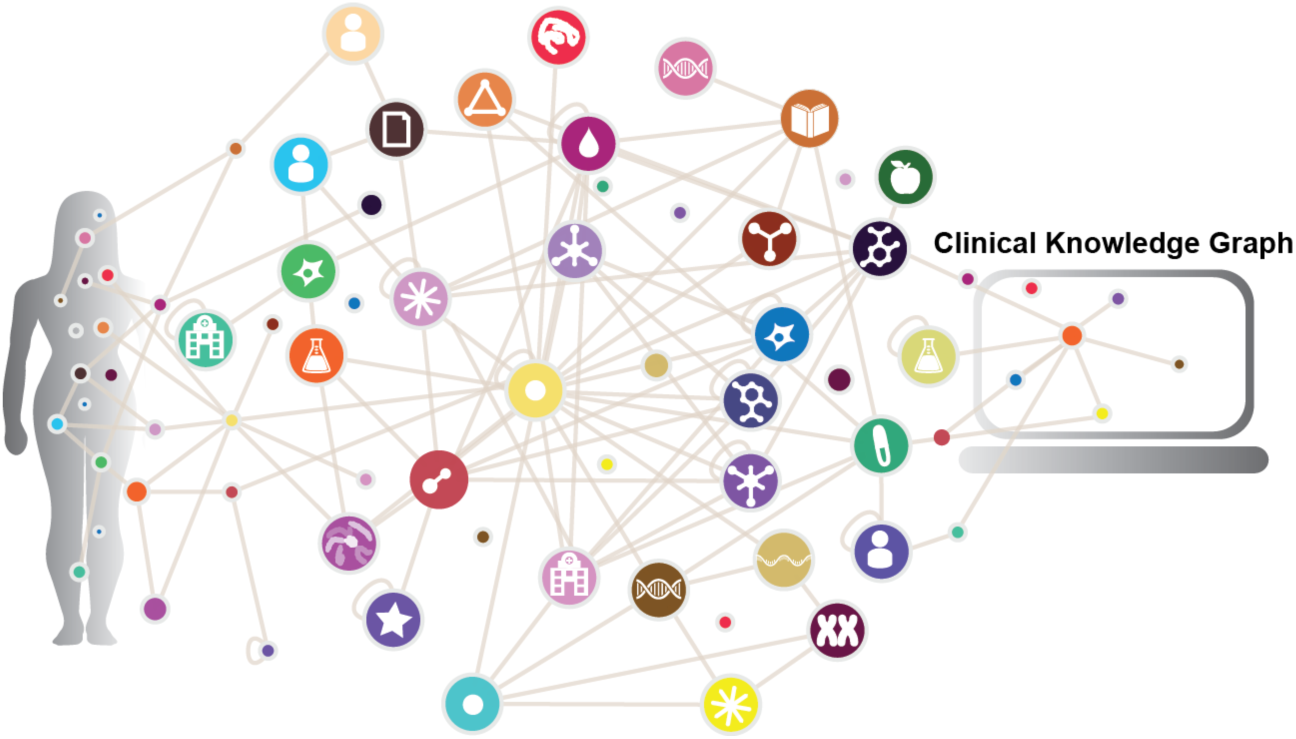

## Introduction

Over the last two decades, the paradigm of evidence-based precision medicine has evolved towards a comprehensive analysis of the disease phenotype. This requires seamless integration of an enormous amount of diverse data, such as clinical, laboratory and imaging data, multiomics data (genomics, transcriptomics, proteomics or metabolomics), and electronic health records (EHRs) (Leopold and Loscalzo, 2018). In our own work, we have recently found that this more fine-grained definition of disease that combines clinical and molecular data, can provide a deeper understanding of individual’s disease phenotype and reveal candidate markers of prognosis and/or treatment (Coscia et al., 2018; Doll et al., 2018, 2019a). Moreover, it is now accepted that mutiomics data aggregation is crucial for generating new hypotheses and developing biomedical knowledge that ultimately translates into clinically actionable results (Lee et al., 2018). The biomedical research community has long recognized the need to collect, organize and structure relevant data resulting in community-wide adoption of multiple biomedical databases (Suppl. Table S1). However, harmonization and integration of the available data is challenging because they are often diverse, heterogeneous and distributed across multiple platforms. Moreover, a significant amount of scientific data and knowledge is only “stored” within millions of unstandardized journal publications that may, to some degree, be captured by text mining efforts (Cook and Jensen, 2019; Pafilis et al., 2013). Thus, the lack of interoperability between the systems and formats used for managing biomedical and clinical data, and challenges associated with the vastness of the data and knowledge, are creating major bottlenecks towards personalized medicine and extracting high-quality, clinically actionable information. This also limits the degree to which results of omics analyses have been integrated into the clinical decision-making process.

Although genomic insights have played a major role in inspiring efforts in personalized medicine, it is increasingly evident that genomics alone cannot explain most disease phenotypes, and that multiomics, in general, and proteomics, in particular, must be taken into consideration (Doll et al., 2019b; Rodriguez and Pennington, 2018; Zhang and Kuster, 2019). Over the last decade, the technology of mass spectrometry MS-based proteomics has advanced greatly and now provides a comprehensive view of biological processes, cellular signaling events and protein interplay (Aebersold and Mann, 2016). Moreover, improvements in sample preparation workflows, MS instrumentation, automation and data analysis software have dramatically enhanced the ability of this technology to more rapidly analyze samples of increasing complexity (cell lines, tissues and body fluids), with increased sensitivity and accuracy (Bache et al., 2018; Bian et al., 2020; Bruderer et al., 2015; Demichev et al., 2020; Gessulat et al., 2019; Heusel et al., 2019; Krieger et al., 2019; Kulak et al., 2014; Meier et al., 2015, 2018; Post et al., 2017; Samaras et al., 2019; Sinitcyn et al., 2018; Slavov, 2020; Tiwary et al., 2019; Zhou et al., 2017). However, currently used MS-based proteomics workflows were conceptualized more than a decade ago, and rapidly increasing data volumes are posing new challenges for the field. An even larger and growing bottleneck in high-throughput proteomics is the difficulty of interpreting the quantitative results obtained, which are typically presented as large spreadsheets, to formulate a biological or a clinical hypothesis. Thus far, only a handful of tools have been aimed at alleviating this problem (Choi et al., 2014; Tyanova et al., 2016), and, as a consequence, proteomics researchers frequently spend an extraordinary amount of time understanding and generating hypotheses from the data, especially when more data types are involved. There is a need for solutions that not only integrate multiple data types, but also capture the relationships between molecular entities at work and the resulting disease phenotype. Moreover, we see an increasing need for more inclusive solutions that provide those with little or no MS-based proteomics expertise with tools for extracting high-quality information from proteomics data in a more user-friendly manner. Therefore, a knowledge-based, knowledge-generating platform that integrates a range of databases and scientific literature information with omics data into an easy to use workflow would empower discovery science and clinical practice.

Networks and graphs have emerged as natural ways of representing connected data also in biology (Barabási and Oltvai, 2004; Barabási et al., 2011; Strogatz, 2001). Efforts during the last decade have organized large amounts of diverse information as collections of nodes (entities) and edges (relationships) especially in user-centric search engine optimization and recommender systems (Balaur et al., 2017; Fabregat et al., 2018a; Himmelstein and Baranzini, 2015; Himmelstein et al., 2017; Mughal et al., 2017). The resulting flexible structure, called a knowledge graph, allows quick adaptation of complex data and connections through relationships. Their inherent inter-connectivity enables the use of network analysis techniques to unveil hidden patterns and infer new knowledge (Yoon et al., 2017). Furthermore, knowledge graphs are computationally efficient and scale to very large sizes as exemplified by social graphs analysis (Fabregat et al., 2018a; Have et al., 2013; Lehmann et al., 2012). Knowledge graphs are already used in the context of health care integrating EHRs into a symptom-disease-therapy matching process (Rotmensch et al., 2017).

Here, we take this concept into a new direction and describe a knowledge graph framework that facilitates harmonization of proteomics with other omics data while integrating the relevant biomedical databases and text extracted from scientific publications. Termed the Clinical Knowledge Graph (CKG), it constitutes a graph database of millions of nodes and relationships. Furthermore, our platform provides clinically meaningful queries and advanced statistical analysis tools, enabling automated data analysis, knowledge mining and visualization. The CKG incorporates community efforts by building on scientific Python libraries (Virtanen et al., 2020), which also makes the platform reliable, maintainable and continuously improving. The entire system is open source and permissively licenced (MIT). It enables repeatable, reproducible and transparent analysis in both standard workflows and interactive, Jupyter Notebook-based exploration. To illustrate the different uses and benefits of this platform, we apply it to a range of published studies, reproducing previous results in drastically shorter time and unveiling additional, actionable knowledge.

## Results

### Overview of Clinical Knowledge Graph (CKG) Architecture

The CKG includes several independent functional modules that serve to: (1) format and analyze proteomics data (analytics_core); (2) construct a graph database by integrating available data from a range of publicly accessible databases, user-conducted experiments, existing ontologies and scientific publications (graphdb_builder); (3) connect and query this graph database (graphdb_connector); and (4) facilitate data visualization, repository and analysis via online reports (report_manager) and Jupyter notebooks (Figure 1A). The latter have become the de-facto standard for accessible, reproducible and sharable data analyses (Perkel, 2018). This architecture allows for a seamless data harmonization and integration, as well as user-supplied analysis. It also facilitates data sharing and visualization, and interpretation based on detailed reports developed to emphasize clinically actionable points. In the next several sections, we will describe individual modules and the knowledge graph construction process in more detail.

**Figure 1.**
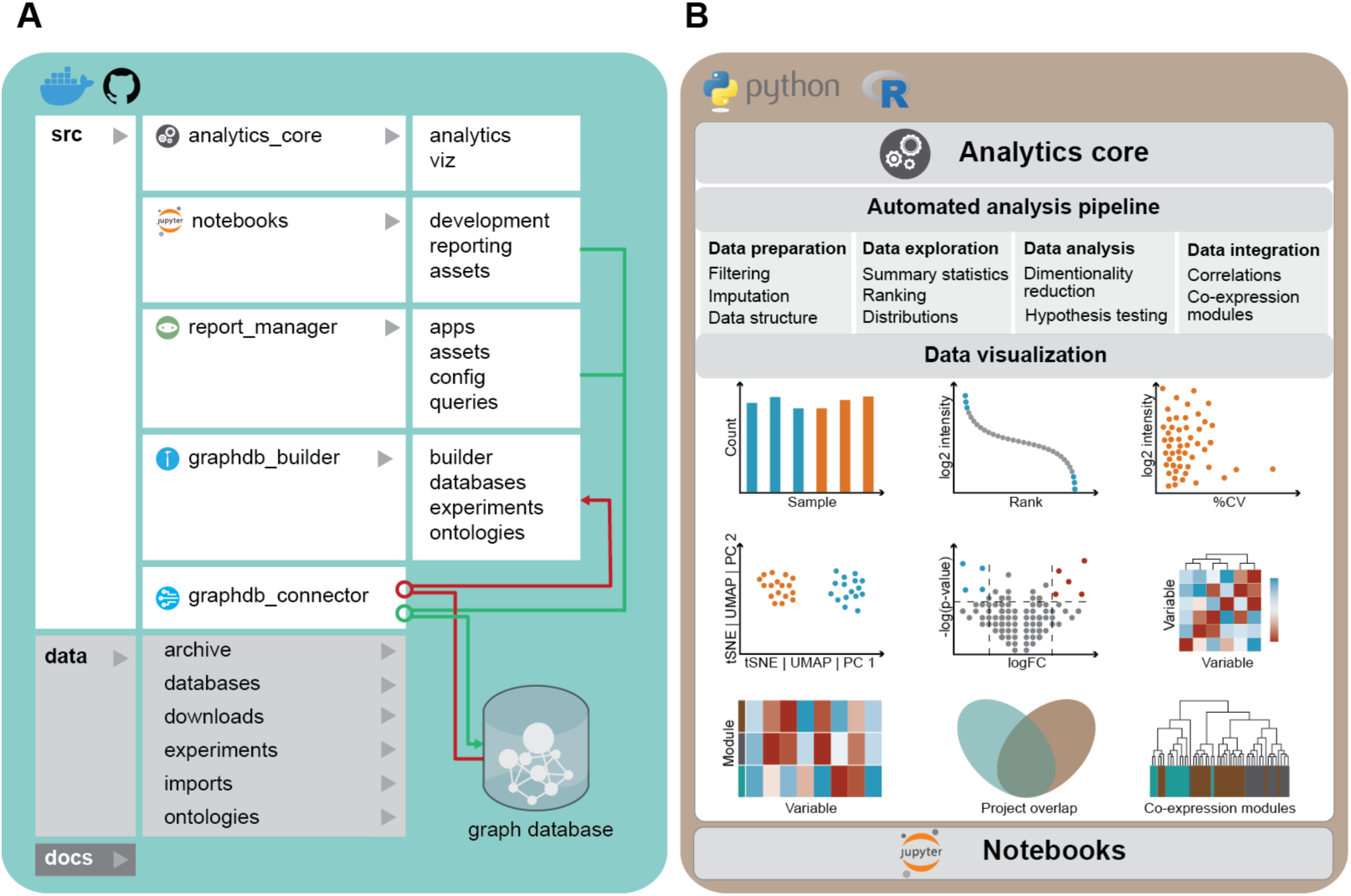
The Clinical Knowledge Graph architecture. A) CKG functionality is implemented in Python and contains several independent functional modules responsible for connecting to the graph database (graphdb_connector), building the graph (graphdb_builder), analysing and visualizing experimental data (analytics_core) and a repository of Jupyter notebooks with analysis templates. The code is accessible at https://github.com/MannLabs/CKG or as a complete Docker container. B) CKG analytics core. This module, developed in Python, makes the analysis transparent and efficient using well tested and up to date algorithms. It centers on statistical and visual data representation and comprises all the main steps in a data science pipeline: data preparation, exploration, analysis, and visualization. Moreover, the modules also allow analysis and integration of other data types, as well as R packages and functions.

### The Analytics Core as an Open Proteomics Analysis Framework

The first step in downstream analysis of proteomics data requires a comprehensive and versatile collection of statistical, machine learning and visualization methods. Tools such as MSstats or Perseus have advanced proteomics by providing multi-purpose statistical and bioinformatics tools for the analysis of quantitative MS-based proteomics data (Choi et al., 2014; Tyanova et al., 2016). MSstats is an R-based package requiring some programming expertise, whereas Perseus is a Windows desktop application aimed at biologists and MS-specialists. Taking these and other efforts as a reference, we developed the Analytical Core that encompasses the desired functionality in a transparent and efficient manner. We chose Python and its associated scientific stack because this allows us to adopt well tested and up to date algorithms while avoiding reimplementing already existing methods. The functionality implemented in the Analytics Core centers on statistical and visual data representation and covers all main areas of computational protemics, such as expression, interaction and post-translational modification-based proteomics (Figure 1).

We designed the Analytics Core to comprise the main steps in a data science pipeline: *data preparation* (filtering, imputation, data formating), *data exploration* (summary statistics, ranking, distributions), *data analysis* (dimensionality reduction, hypothesis testing, correlations) and visualization. The Analytics Core goes beyond previous efforts by integrating other data types in addition to proteomics (i.e. clinical data, multiomics, biological context and text mining). Furthermore, to complement the extensive Python portfolio, we incorporated functions optimized in the R language (i.e. SAMR and WGCNA (Langfelder and Horvath, 2008; Pei et al., 2017; Tusher et al., 2001))(Suppl Table S2).

We included a visualization module (viz) that covers both basic plots (e.g. barplot or scatterplot) and more complex ones (e.g. network, Sankey or polar plots). We implemented these visualizations with Plot.ly (https://plot.ly/), a graphing library compatible with multiple programming languages (i.e. Python and R). This way, visualizations created by the CKG framework and stored for instance in JavaScript Object Notation (JSON) can be easily exported and used from other languages (see Suppl. Methods).

The initial *data preparation* step structures the quantified measurements (filtering, imputation, formatting, normalization), starting with filtering out proteins identified only in few of the samples (Suppl Table S2). This filtering step can be specified as a maximum percentage of missing values (default) or as a minimum number of values present per condition (group) or in the entire dataset. For imputation, we implemented several methods that account for missing values of different nature, including he k-Nearest Neighbors imputation method (KNN) which assumes that the values are Missing Completely At Random (MCAR), and Probabilistic Minimum Imputation approach (MinProb) for missing values that are considered Missing Not At Random (MNAR)(default) (Lazar et al., 2016). These two methods can also be combined in a mixed imputation method that considers the percentage of missing values in order to assume missingness due to MCAR (i.e. missingnes <50%) or MNAR otherwise, and applies KNN or MinProb, respectively. This step results in a complete matrix called the “processed data frame” and forms the basis for downstream analysis.

Next, we implemented the *data exploration* step into the workflow to collect summary statistics from the original data (number of proteins, peptides, etc.). Additionally, it ranks identified proteins according to their average quantified intensity (Label-Free Quantification, LFQ (Nahnsen et al., 2013)) and calculates protein coefficients of variation, which can serve as a quality metric.

The subsequent *data analysis* part includes a dimensionality reduction step and enable visualization of the high dimensional proteomic datasets using two or three-dimensional representations. We implemented linear dimensionality reduction (Principal Component Analysis (PCA) (default) (Hotelling, 1933)), and non-linear approaches (t-distributed stochastic neighbor embedding (tSNE) (Van Der Maaten, 2009)) and Uniform Manifold Approximation and Projection (UMAP) (McInnes et al., 2018)).

The Analytics Core enables hypothesis testing, particularly methods for identifying proteins changing significantly between conditions (groups). The default method is Analysis of variation (ANOVA), but others, such as ANOVA for repeated measurements (ANOVA-rm), t-test (independent or paired) or Significance Analysis of Microarrays (SAM), are also available (Tusher et al., 2001). By default, the analytics core identifies the appropriate test based on the experimental design (e.g. independent *vs* paired, ANOVA *vs* ANOVA-rm). We also implemented several methods to correct for multiple hypothesis testing such as Benjamini-Hochberg False Discovery Rate (BH-FDR) (default) (Benjamini and Hochberg, 1995), or permutation-based FDR, which is used only if the number of permutations specified (default set to 250) is sufficiently large to avoid overestimating false positives.

Strategies for global protein-protein correlation analysis include as default Pearson correlation analysis corrected for multiple testing, which returns a network with identified clusters of correlating proteins (Louvain clustering method). Further, functional enrichment analysis (Gene Ontology and Pathways (Ashburner et al., 2000; Fabregat et al., 2018b)) enables extraction of potential hypothesis-generating information regarding the functional consequences of proteomics perturbation as an ultimate step in the proteomics analysis (Suppl Table S2).

Due to its modular design, the analytics core functionality can be used within the CKG framework but also independently by importing it from Python. Similarly, the analysis and visualization functionalities are not limited to proteomics data but can handle any type of data in matrix format. The open design promotes easy integration of new analysis methods and visualizations. As an example, the above-mentioned UMAP dimensionality reduction was readily imported through the corresponding library (McInnes et al., 2018). Our integration of Jupyter Notebooks, an open source tool that allows mixing of text, graphics, code and data in a single document (Lamb et al., 2006; Mendez et al., 2019; Perkel, 2018; Rule et al., 2019) enables standard or bespoke analysis pipelines, including addition of existing or user-implemented functionality from the Python or R ecosystems.

### Building and Populating a Graph Database

To achieve seamless annotation and integration of proteomics data with other types of disparate information such as the results of other omics experiments and/or literature information, we constructed a graph database that would inherently connect large and heterogeneous data. We chose the open source Neo4J database platform as our current backend because of its performance, industry acceptance and associated Cypher query language (https://neo4j.com/). To build the knowledge graph, we first wrote a library of parsers (graphdb_builder) with associated configurations for each ontology, database, and type of experiment. They download the data from online resources, extract information and generate entities (nodes) and relationships, both of which can have properties (attributes), such as name or description in protein nodes. The parsers use paired configuration files that specify how ontologies, databases or experiments need to be interpreted. This design allows unrestricted integration of new resources or processing tools, as well as easy adaptation to changes in the input format. Output formats tend to change from time to time but this only affects one parser/configuration that can be easily adapted. For example, the current CKG’s proteomics parser accepts output data from data dependent or data independent as well as isobaric labeling experiments from commonly used programs such as MaxQuant or Spectronaut (Bruderer et al., 2015; Cox and Mann, 2008), but this can easily be modified for additional data outputs as new processing programs emerge.

Once the ontology, database and experiment files are standardized, formatted and imported, the graphdb_builder module loads them into the graph database with a set of Cypher queries that create the corresponding nodes and relationships. To illustrate, disease nodes are identified through Disease Ontology identifiers (e.g. DOID:150) (Schriml and Mitraka, 2015), tissue nodes use Brenda Tissue Ontology (e.g. BTO:0000759) (Gremse et al., 2011; Jeske et al., 2019) whereas proteins follow UniProt codes (e.g. P01116) (UniProt Consortium, 2018). Relationships are defined by source (‘start node’) and target (‘end node’) and type of association (‘type’). Relationship types can be used to query for all the source/target node pairs linked by a specific type. The builder.py module simplifies the update of the CKG database (Methods).

The data model we built connects 33 different node labels with 51 different relationship types (Figure 2A). It enables predefined queries about experimentally determined protein hits, regarding their association to the diseases studied (ontological associations), drugs or annotated gene ontology terms and pathways. These types of queries could result in insights into altered functions, suggest drugs for regulated proteins and even connections to metabolites and diet (food) to reveal possible confounding factors. Moreover, new queries can readily be constructed with some background in Cypher, the query language used by Neo4j and other graph databases. Cypher syntactically represents the structure of the graph with nodes enclosed within parentheses and relationships depicted with arrows between the nodes (Figure 2B).

**Figure 2.**
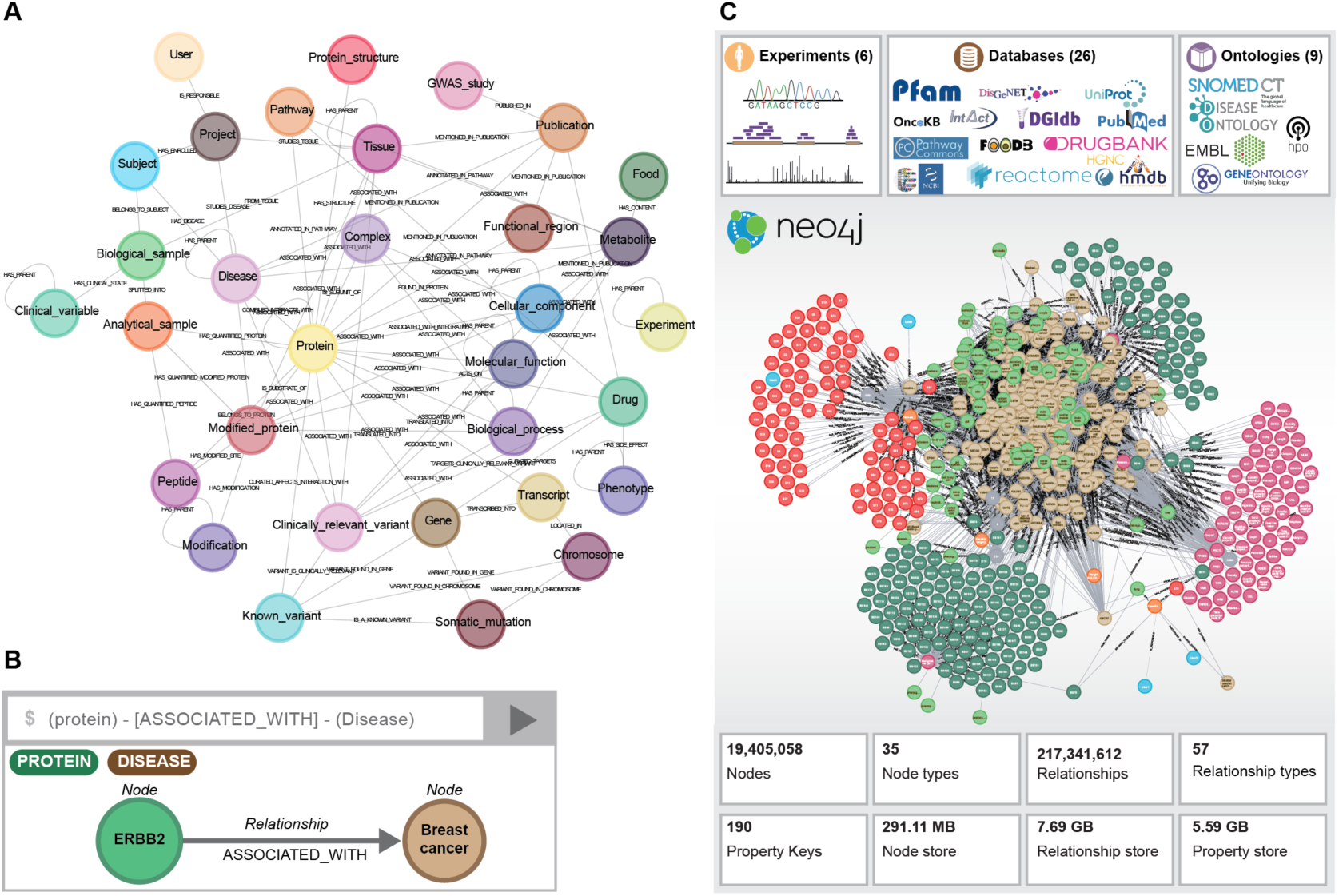
A knowledge graph of millions of nodes and relationships. A) Data model: Node labels (Protein, Metabolite, Disease, etc) and relationship types connecting them (HAS_PARENT, HAS_QUANTIFIED_PROTEIN, etc). The nodes and relationships have been designed to answer biomedical questions including those relevant for the clinic. B) Cypher query example. C) As depicted, CKG integrated 6 experimental studies, 9 ontologies and 26 databases into a knowledge graph with more than 16 million nodes and more than 220 million relationships.

To make experimental proteomics data FAIR (findable, accessible, interoperable and reusable (Wilkinson et al., 2016) we designed a data model structure capable of supporting storage of standardized metadata (studied disease, interventions, etc.) around each research project, defining unique identifiers for enrolled subjects, collected biological samples and analyzed samples (Suppl. Figure S1A). In this graph structure, clinical data is integrated by building relationships (HAS_QUANTIFIED_CLINICAL) between biological samples and clinical variables codified using the Systematized Nomenclature of Medicine Clinical Terms (SNOMED-CT) (Millar, 2016), while proteomics data is used to create new relationships between analytical samples and proteins (HAS_QUANTIFIED_PROTEIN). In both cases, quantified values are stored as relationship properties.

### Clinical Knowledge Graph Includes Millions of Nodes and Relationships

The CKG database is constantly growing but currently collects annotations from 26 biomedical databases using 9 ontologies, and organizes this information into 16 million nodes connected by 220 million relationships (Figure 2C). We purposfully built in some redundancy by including biomedical databases that provide the same type of relationships (i.e DISEASES (Pletscher-Frankild et al., 2015) and DisGeNET (Piñero et al., 2019)), which we used to assess overlap and disagreement of sources (Suppl. Figure S2A).

More than 50 million relationships involve ‘Publication’ nodes (MENTIONED_IN_PUBLICATION) linking scientific publications, coded with Pubmed identifiers, to proteins, drugs, diseases, functional regions and tissues (Suppl. Figure S2B). They were derived from text mining of almost 7 million abstracts and full-text articles (Cook and Jensen, 2019; Pafilis et al., 2013), thus encapsulating aspects of the accumulated biomedical knowledge in peer-reviewed publications.

We found the graph structure to be easily scalable, facilitating the integration of new ontologies, databases and experiments. The number of nodes and relationships can increase without appreciably affecting the performance of graph databases (Fabregat et al., 2018a; Have et al., 2013). Furthermore, this inherent flexibility allows nodes and relationships originally designed to provide biomedical context primarily for large-scale proteomics data interpretation to be readily remodeled to integrate other omics datasets. For instance, to integrate metabolomics data we can link metabolite nodes already present in the database to analytical samples (HAS_QUANTIFIED_METABOLITE). The existing relationships connecting metabolites to nodes such as pathways, proteins, food, tissue or disease, would aid interpretation and seamless integration with proteomics results (Zhang and Kuster, 2019).

The CKG framework provides an infrastructure that facilitates exploiting the existing connections in the graph and the already implemented and optimized graph algorithms in Neo4j and Python libraries (NetworkX) (Hagberg hagberg et al., 2008). For example, when a new project is integrated the default analysis identifies similar projects in the graph and the results of these comparisons are shown in the project report. This functionality compares projects based on the overlap of identified proteins (Jaccard and overlap similarity) or similar protein profiles (cosine similarity). In addition, these algorithms can be used to build new links between nodes based on similarity of their connected subgraphs, a strategy known in graph learning as link prediction or graph completion (Liben-Nowell and Kleinberg, 2004; Nickel et al., 2015). For instance, we used this functionality to map Gene Ontology biological processes to metabolic pathways (Suppl. Table S3). This helps to better interpret functional enrichment results or to connect currently disconnected nodes and extend their annotations, i.e. (Biological_processes-[:ASSOCIATED_WITH]-(Metabolite). Graph-based predictions have been used in multiple scenarios including drug repurposing, protein-protein interaction (PPI) prediction, disease comorbidity risks or diet-based cancer therapy associations (Cheng et al., 2018; Halu et al., 2019; Menche et al., 2015; Veselkov et al., 2019). All the types of relationships mined in those studies are part of the CKG and can repeatedly be modelled in the same manner every time new data is integrated. Moreover, besides the database functionality offered by Neo4j, CKG also provides a library of optimized graph algorithms that run within the database framework. These algorithms efficiently implement graph analysis tools such as path finding, centrality measurements, community detection or similarity functions among others. All these algorithms are directly available in the CKG to effectively identify hidden patterns and generate predictions based on the connected data. Lastly, we developed CKG to enable user interaction either through a web interface or desktop application, or programmatically through a bolt protocol, in all cases through (pre-packaged) Cypher queries (Methods).

### A Framework to Extract Actionable Knowledge

A main goal of the CKG is to combine the power of the analytics module with the massive prior information integrated into the graph database to best interpret MS-based proteomics or other omics experiments. Harmonization of these heterogeneous but connected data sources enables the implementation of standard analysis pipelines that report results immediately, replacing weeks of manual work in a more consistent format. These standard reports provide initial evaluation of the quality of the generated data, highlight relevant hits, and contextualize these hits in relation to different biomedical components in the graph. Standard analysis workflows can be defined for different data types (for instance, clinical, deep sequencing and proteomics data). The report manager component (report_manager) orchestrates the creation and updates experimental projects and the automatic analysis, visualization and knowledge extraction (Figure 3).

**Figure 3.**
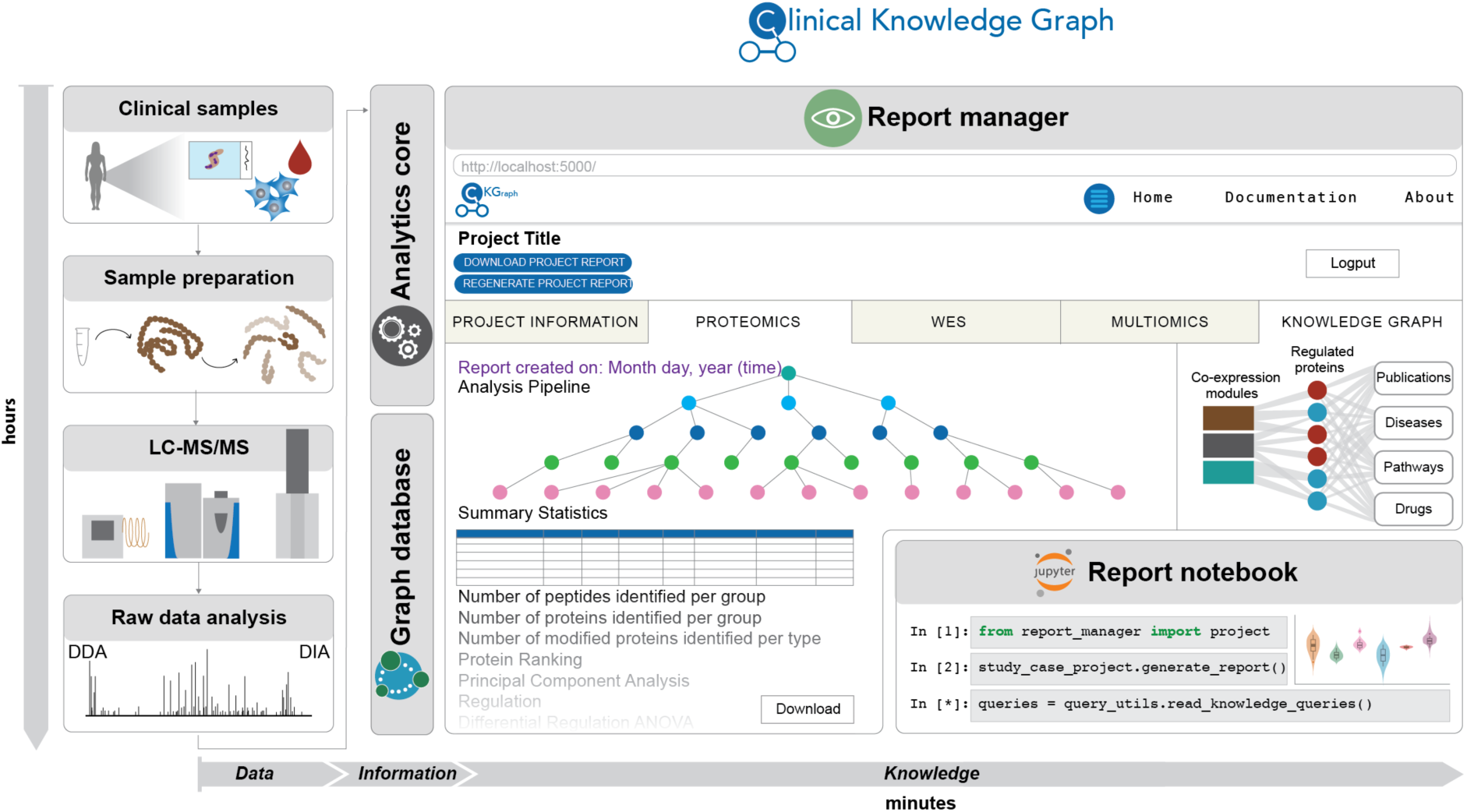
The report_manager coordinates multiple processes. This CKG module includes a collection of dashboard applications that interface with the database and display statistics, create and upload new projects, and run the automated analysis pipelines. The resulting report includes multiple tabs, one for each data type analysed, and a multiomics tab when multiple data types are analysed together, and a knowledge graph that summarizes the results obtained in the previous tabs. This report can be viewed and downloaded through the Dash application in the browser, and accessed through a Jupyter Notebook.

The report manager was implemented as a collection of dashboard applications that interface with the database for an overview of the knowledge graph (Home) (Materials), to create and upload new clinical proteomics projects (Project creation and Data upload) and to run automated analysis pipelines (ProjectApp). This defines a workflow from project idea to knowledge-based analysis report (Suppl. Figure S1B). The project creation and data upload step generate the nodes and unique identifiers in CKG (internal identifiers) for the project (P), the enrolled cohort subjects (S), biological samples collected (BS) and analytical samples analyzed by MS-based proteomics (AS). This includes description, starting and ending dates, and experimental design as well as diseases, tissues and clinical interventions (Suppl. Figure S1C), which provides prior knowledge, such as known proteins associated with these diseases and tissues or literature related to them. This information can also identify similar projects in CKG, or to estimate the expected statistical power including required effect sizes. Once the clinical and/or proteomics data are ready and processed, they are integrated into the graph by the ‘Data upload’ dashboard app (Suppl. Figure S1D).

Uploading the data triggers the builder’s import and load process and generates all the necessary relationships within the new project, including the links to proteins, peptides and protein modifications quantified in the project. The new links are used by the report manager module to analyze the different data types relevant for the project. The analyses are predefined using configuration files that break down the steps of the complete analysis workflow in a standardized, flexible and scalable manner. They describe the input data, the analysis to be performed and the parameters to be used, and how the results are visualized. The final report is then composed of the sequence of created visualizations for each data type divided into different tabs in the dashboard (plots and tables). When a project contains multiple data types, CKG allows studying different types in combination, for instance global correlation or WGCNA analysis of clinical and proteomics data. To facilitate interpretation of all these results, the Knowledge Graph tab summarizes all the analysis’ outputs together with connective knowledge extracted from the graph (see case studies below).

Apart from being viewable in the browser, all the reports, analysis results and visualizations can be downloaded as a single compressed file containing tables and figures in ready to publish format. Moreover, they are also available in Hierarchical Data Format (HDF5), a standard and scalable file format supported by many programming languages, enabling interoperability. This design facilitates continuous integration of newly developed analyses and visualizations (Suppl. Figure S3). Furthermore, configuration files can be shared, which fosters transparency and facilitates replicability and reproducibility.

### Automated CKG Analysis for Liver Disease Biomarker Discovery

To illustrate how CKG accelerates and extends both the analysis of the data and its interpretation, we use its default pipeline on a proteomics study of Non-Alcoholic Fatty Liver Disease (NAFLD), in which we had confirmed several known markers and revealed new ones associated with liver disease progression such as ALDOB, APOM, LGALS3BP, PIGR, VTN, and AFM (Niu et al., 2019). Given only standard configuration files describing the sequences of analysis and visualizations steps: *data preparation*, *data exploration* and *data analysis*, we asked how comprehensive the CKG interpreted the data compared to our previous manual analysis (Figure 4).

**Figure 4.**
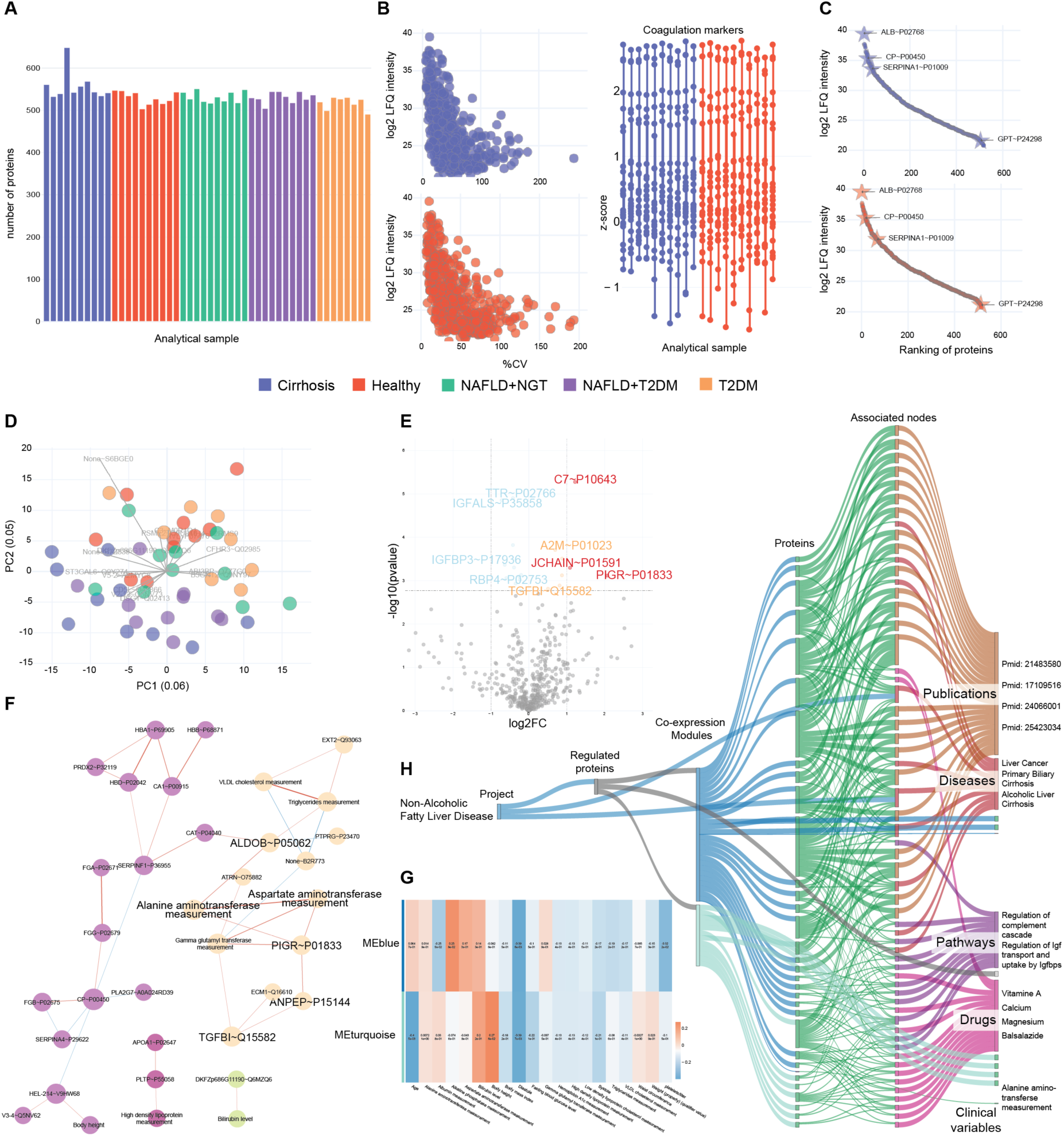
CKG automated default analysis of the Non-Alcoholic Fatty Liver Disease Study. CKG’s automated analysis pipeline reproduced the results (Niu et al., 2019). The visualizations were generated by the report manager and downloaded from the dashboard app. The plots correspond to the analyses performed on the proteomics data (Proteomics tab; A-E), the WGCNA and Clinical-Proteomics correlation analyses (Multiomics tab; F-G) and the summary tab (Knowledge graph; H).

In the default analysis pipeline, each data type collected in the project (clinical, proteomics) was analyzed automatically and independently. For the clinical data, the CKG summarized clinical characteristics of the cohort and highlighted variables that have significantly different variances between the studied groups (Healthy, Normal Glucose Tolerance (NGT), Type 2 Diabetes (T2D), NGT with NAFLD, T2D with NAFLD and Cirrhosis). These analyses confirmed the expected significant differences between groups in the levels of the liver enzymes Alanine aminotransferase (ALAT), Aspartate aminotransferase (ASAT) and Alkaline phosphatase measurement (ALP), or Hemoglobin A1c among others. Likewise, pairwise Pearson correlation analysis identified associations between clinical parameters such as the levels of liver enzymes (ALAT/ASAT/ALP) and systolic and diastolic measurements and the studied groups.

The proteomics default analysis started with the *data exploration* step. This yielded an overview of the identified peptides, proteins and protein modifications as well as a detailed summary of descriptive statistics of the proteomics data matrix (mean quantification values, standard deviation, quartiles, etc.) (Figure 4A). This *original dataframe* can be used to evaluate the quality of the data and identify significant outliers. CKG continued with a visualization of proteome coverage, dynamic range, protein coefficients of variation (CVs) among samples and when available sample quality control based on known tissue quality markers (Figure 4B) (Geyer et al., 2019).

These rankings were automatically annotated with curated information mined from the knowledge graph – in this case highlighting already known markers linked to the studied disease or related ones (NAFLD, cirrhosis etc.). This was implemented as a simple Cypher query: ((:Protein)-[:IS_BIOMARKER_OF_DISEASE]-(:Disease)). In our present study, the CKG marked known liver disease markers on the abundance ranking plot such as Albumin (ALB), Ceruloplasmin precursor (CP), Alpha-1-antitrypsin (SERPINA1) and Alanine aminotransferase 1 (GPT) in the different groups (Figure 4C).

Next, the default *data analysis* used PCA to perform feature reduction on the data (*processed* data frame) (alternatively, t-SNE or UMAP) (Figure 4D). Analysis of variance (ANOVA) with posthoc tests then determined statistically significant differences across all studied groups and between particular pairs of groups after adjusting for multiple hypothesis testing. This resulted in a comprehensive data table that also provides alternative effect size measurements such as Hedges ratio (Hedges and Olkin, 1985). Posthoc tests are presented as interactive volcano plots, with information about up- and down-regulated proteins with a predefined significance threshold (i.e. fold-change > 2 and FDR < 0.05) (Figure 4E). The CKG automatically reproduced all our previous manually analyzed results showing dysregulation of proteins involved in immune system regulation and inflammation, such as Complement component C7 (C7), Immunoglobulin J chain (JCHAIN), Polymeric immunoglobulin receptor (PIGR) and alpha‐2 macroglobulin (A2M), an established clinical marker of liver fibrosis for which CKG reported 14 publications establishing this connection. In addition, CKG highlighted the TTR-RBP complex (TTR and RBP4) as downregulated in cirrhotic patients compared to healthy ones. This complex is involved in the retinoid metabolism and its dysregulation has been previously linked to hepatic diseases and alteration of the extracellular matrix deposition leading to fibrosis (Shirakami et al., 2012). Further, the report revealed literature and database associations between the regulatory role of CD5 antigen-like protein (CD5L) and cirrhosis, hepatocellular carcinoma and other liver diseases (Sarvari et al., 2013). These metabolically interesting findings were missed in our manual analyses but were prioritized by CKG’s automated pipeline, which maximizes the information extracted from significantly regulated proteins in the different conditions. To visualize correlated protein changes as a network, the default analysis connected proteins with significant associations (Pearson correlation coefficient > 0.5; FDR<0.05). On this graph CKG used the Louvain algorithm to detect communities or clusters of highly correlating proteins (Blondel et al., 2008). This revealed potentially clinically relevant connections such as the association of a cluster composed by PIGR, and DPP4 and TGFBI to liver fibrosis. In addition to the correlation network, the CKG also used its background knowledge of millions of protein interactions. Here, the PPI network of significantly regulated proteins revealed six main clusters grouping extracellular matrix remodelers, complementary components and inflammation markers, while also connecting two of the candidate biomarkers (PIGR and JCHAIN).

The last part of the default data analysis pipeline associated the differentially regulated proteins to drugs, diseases and publications, and to enriched biological processes and pathways. This unveiled additional dysregulated pathways in NAFLD overlooked in our previous analysis, including ‘regulation of Insulin-Like Growth Factor (IGF-1) transport’ and ‘uptake by Insulin-Like Growth factor Binding Proteins (IGFBPs)’, which were linked to changes in IGFBP3, acid-labile subunit (IGFALS). Notably, associations between IGFBPs and insulin resistance in NAFLD have recently been reported and investigated for causal relationships and therapeutic potential (Ahrens et al., 2013; Wittenbecher et al., 2019). CKG also reported protein-disease associations indicating possible disease comorbidities or common disease mechanisms as reflected in their shared dysregulated protein signatures, such as primary biliary cirrhosis, liver cancer, hepatitis, biliary tract disease, and pancreas disease.

The presence of various data types in the project next triggered the default multiomics analysis pipeline. Here, CKG performed a global clinical-proteomics correlation analysis (Wewer Albrechtsen et al., 2018) (Figure 4F). In this case clinical liver enzyme values clustered together with HbA1c, fasting glucose levels and several candidate biomarkers of liver fibrosis and cirrhosis (PIGR, TGFBI, ANPEP, C7, etc.). The CKG also used WGCNA to obtain modules of co-expressed proteins instead of individual proteins that are related to clinical variables (Figure 4G).

Finally, the automated analysis pipeline summarized all the results of the clinical, proteomics and multiomics analyses. Represented in a Sankey plot (Schmidt, 2008), it connected co-expression modules, differentially or co-regulated proteins and clinical variables, related diseases, pathways, drugs or publications co-mentioning the disease and the regulated proteins (Figure 4H). This visually summarized the entire multi-analysis, thereby helping to understand the underlying molecular mechanisms of the disease or disease comorbidities.

Importantly, the entire default pipeline took less than five minutes and captured basically all of the insights gleaned from our previous manual analysis, which had taken weeks. The advantage of using the CKG became particularly clear in the interpretation of the differentialy abundant proteins, which had comprised extremely time-consuming literature and database searches for known/published protein-disease associations and knowledge gathering (Niu et al., 2019). In summary, apart from drastically speeding up the data analysis pipeline and capturing biological findings in a more comprehensive manner, CKG added several novel leads to our previous manual analysis.

### CKG Enables Multi-proteomics Data Integration for Cancer Biomarker Discovery and Validation

The CKG is built to easily and quickly integrate different proteomics data types (i.e. interactomics, postranslational modifications) and contains analytical tools to extract relevant information from these data types and to contextualize the findings with knowledge from the graph. To explore these capabilities, we reanalyzed a recent study in which we identified Cancer Testis Antigen family 45 as a biomarker for long-term survival in ovarian serus adenocarcinoma and described its mode of action (Coscia et al., 2018).

Multi-dimensional proteomics, phosphoproteomics and interactomics were modeled as different connected projects in CKG and analysed independently using the default analysis adapted to each data type (proteomics, interactomics and phosphoproteomics). The proteomics analysis followed the default pipeline already described, which reproduced in an automated manner the published results identifying CT45 as significantly more highly expressed in patients with long-term remission after chemotherapy (Figure 5A). CKG performed Kaplan-Meier survival analysis of high (top 25%) vs low CT45 expression, confirming that the former had significantly longer disease-free survival (Figure 5B). CKG also confirmed that there was virtually no previous knowledge about cellular roles and functions of CT45. However, CKG’s default analysis immediately produced 24 potential interactors of CT45, with four of them belonging to the PP4 complex and contributed by a human interaction map (Hein et al., 2015). Further interactomics experiments involving reciprocal tagging confirmed these interactions after analysis in the CKG (Figure 5C).

**Figure 5.**
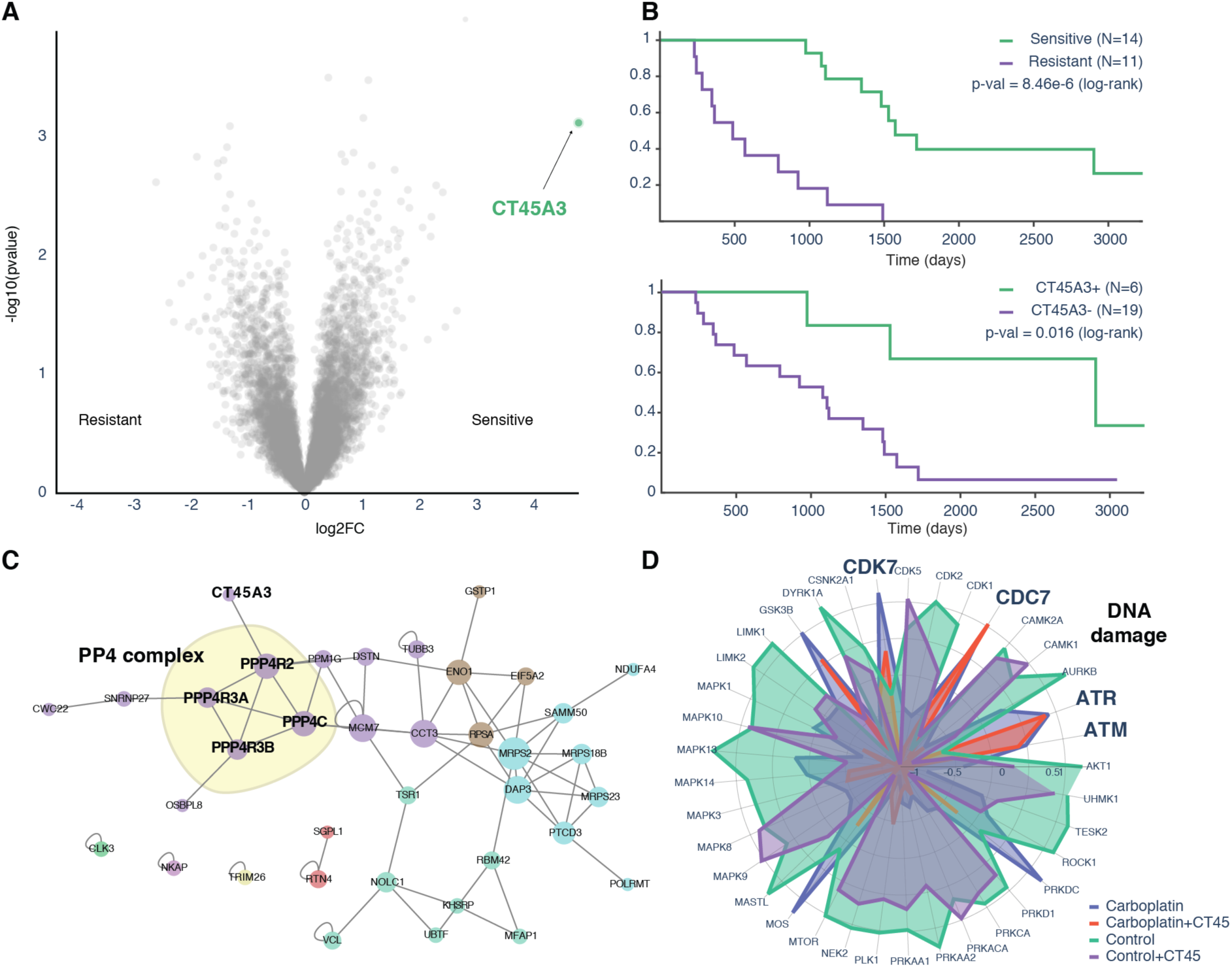
CKG analysis establishing CT45 as a Chemosensitivity Mediator in Ovarian Cancer. A) CT45A3 is highlighted as the only protein significantly regulated when comparing ovarian tumor tissue from chemoresistant and sensitive patients (n=25) (data from (Coscia et al., 2018)). B) CKG’s analysis pipeline estimates the survival function for the clinical groups sensitive and resistant and for patients with different CT45A3 protein signatures (top 25% expression CT45+, remaining 75% CT45-). C) Interaction proteomics revealed subunits of the PP4 phosphatase complex as direct interactors of CT45 shown by CKG as cluster in the PPI network confirming known interactors and highlighting potential novel ones. D) Phosphoproteomic analysis in CKG identified significantly regulated sites and linked them to upstream kinase regulators.

The PP4 complex has been linked to DNA damage repair (Gingras et al., 2005), which – together with the fact that patients had undergone chemotherapy inducing DNA interstrand crosslinks - prompted us to investigate the phosphoproteome in cell line models (Coscia et al., 2018). CKG’s detault analyis of the signaling response of CT45 expressing cells vs. controls indeed revealed activation of the relevant DNA damage pathways. Further analysis of the kinases responsible for this phenotype and the connection to other diseases in the CKG associated these sites to upstream kinases, helping to better understand the signaling pathways involved. This pinpointed several known DNA damage kinases (i.e. ATM/ATR) and their correponding regulated substrates (SMC1A_S966/S957,NBN_S615 and PBRM1_S948) providing deeper insights into the mechanisms of action of carboplatin and its negative regulation of proliferation through DDR. Additionally, CKG exposed several other relevant kinases and associations that had eluded manual analysis such as site-specific activation of MAPK activity, as well as differences in CDC7 and CDK7 substrate regulation (Figure 5D). Thus the CKG seamlessly intergrated diffiferent layers of prior data and different dimensions of proteomic data. Although not relevant to this example, the CKG includes similar capabilities for genomic and transcriptomics data, as well as other omics data types, thus allowing further integration.

### Using CKG to Inform Prioritization of Treatment Options for Chemorefractory Cases

After standard treatment options have been exhausted in end-stage cancer, molecular profiling may still reveal druggable targets and opportunities for drug repurposing (ICGC/TCGA Pan-Cancer Analysis of Whole Genomes Consortium, 2020; Pushpakom et al., 2018). Our group has previously used proteomics profiling of cancer tissue to identify alternative therapeutic options by proposing targeted strategies (Doll et al., 2018, 2019b). These kinds of clinically relevant scenarios have shaped the CKG and underly the type of nodes and relationships that we can draw on to efficiently support treatment recommendations. In particular, CKG currently mines more than 350,000 connections between proteins and small molecule compounds targeting them - the approved or investigational drugs (Suppl. Table S4).

In our previous proteomic study of a chemorefractory metastatic case of urachal carcinoma, we proposed lysine‐specific histone demethylase 1 (LSD1/KDM1A) as a possible druggable target (Doll et al., 2018). Here, we extended that study with a broadened default analysis, supplemented with Jupyter notebooks that implement repurposing based on prior knowledge in a reusable pipeline that can be applied in other studies (Figure 6). As we found seveal hundred significantly regulated proteins when comparing lung tumor to non-cancerous appearing tissue a strategy for knowledge-derived prioritization, such as text mining, disease and drug associations became necessary (Corsello et al., 2020; Nowak-Sliwinska et al., 2019; Pushpakom et al., 2018). The CKG combines these approaches by mining the graph to identify drug-target-disease triplets co-mentioned in the literature (3.3 million publications mentioning triplets), to enumerate side effects associated with drugs (72,000 associations), to find similar drugs based on side effects, indications, targets, and to connect drugs with functional pathways.

**Figure 6.**
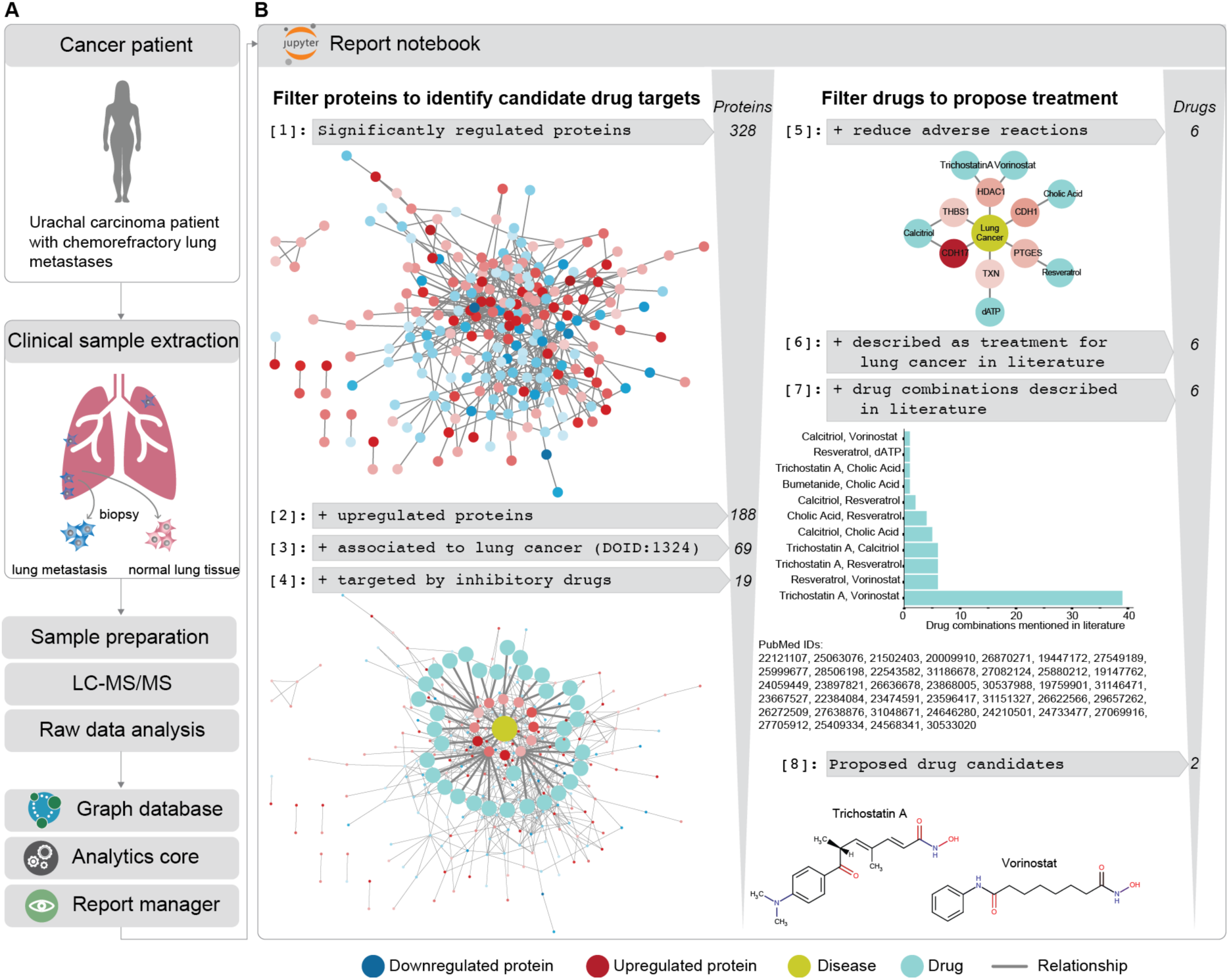
Accelerated prioritization of candidate treatments for a chemorefractory cancer patient. The pipeline mined the CKG database to identify upregulated proteins known to be linked to the studied disease, found inhibitory drugs for these proteins, retrieving reported side effects and ultimately identified possible combinations of the prioritized drugs based on co-mentioning in scientific literature.

We implemented this analysis pipeline in a Jupyter Notebook, which proposed alternative treatments based on the proteomics data and the knowledge extracted from the CKG connections (Figure 6). Running the default analysis resulted in 328 differentially regulated proteins, 188 of which were upregulated (paired t-test, Suppl Table S2). Focusing on proteins with a known association to Lung Cancer (DOID:1324) yielded 69 proteins. Next, the CKG identified drugs that have an inhibitory mechanism for these proteins. One of these targets is LSD1/KDM1A, which was also reported in our original manuscript. However, here CKG not only automatically connected LSD1/KDM1A to tranylcypromine, the drug approved by the tumor board for our patient, but also indicated that trans-2-phenylcyclopropylamine, a small molecule with known potent inhibitory effects on the demethylase, may represent another treatment option (Mimasu et al., 2010). In total, the analysis identified 60 potential drugs targeting 19 of the prioritized proteins. To avoid previously observed adverse reactions, we retrieved all the side effects associated with these inhibitors when used in chemotherapeutic regimens. The pipeline further ranked the remaining drugs according to dissimilar side effects (Jaccard index). Setting a conservative cutoff (<0.2) further prioritized the list, resulting in 6 drugs (Cholic acid, dATP, Resveratrol, Calcitriol, Vorinostat and Trichostatin A) targeting 6 proteins (HDAC1, THBS1, CDH1, PTGES and TXN).

Additionally, the CKG suggested a list of publications that co-mention these drugs with their protein targets and the disease and affected tissue, which in this case highlighted the combination of Vorinostat and Trichostatin A in more than 30 publications. These drugs inhibit HDAC1, a histone deacetylase that induces epigenetic repression linked to tumor progression. Combinations of such inhibitors have been used successfully to inhibit epigenetic silencing and its malignant effects (Rafehi and El-Osta, 2016; Vashishta and Hetman, 2014; Wang et al., 2013). Furthermore, primed by the involvement of both HDAC1 and LSD1/KDM1A in histone modifications, we extended the analysis to find possible connections between these proteins. CKG revealed that they are subunits of several multiprotein complexes. This includes the CoREST complex, which has recently attracted therapeutic interest and for which CKG retrieved a paper describing the inhibition of both HDAC1 and LSD1 (Kalin et al., 2018).

## Discussion

Technological advances are facilitating the generation of increasingly larger omics data sets including those from MS-based proteomics. Despite many promising improvements in bioinformatics tools, there is an ever-larger bottleneck in the analysis, interpretation and especially the extraction of actionable knowledge from these data sets. The lists of differentially expressed proteins generated by existing software tools are still difficult to review manually, especially for non-experts. Bioinformatic results typically provide little guidance for subsequent time-consuming literature searches and integration with existing biological knowledge about diseases, therapies, biological entities and their functions, which is scattered across multiple resources and may be buried in the vast scientific literature. This is especially problematic in clinical studies.

To begin to address these issues, we developed an open framework called the Clinical Knowledge Graph, which analyzes proteomics data with state-of-the-art methods. It automatically annotates omics results with biological and biomedical context extracted from the collective knowledge that has already been built up by the community. To this end, CKG represents prior knowledge, experimental data, and clinical deidentified patient information in a large network. The CKG database harmonizes proteomics data with all this information using a graph structure that naturally provides immediate connections to the identified proteins. Its automated, instantaneous and iterative nature helps in revealing pertinent biological context for better understanding and generation of new hypotheses. Futhermore, the graph structure provides a flexible data model that we have found to be easily extendable to new nodes and relationships. For instance, we were able to quickly expand the library of parsers in the graphdb_builder to connect new ontologies, databases and omics data types, such as imaging data. While the CKG was particularly designed for answering clinically relevant questions, it is equally applicable to other organisms and to any biological study.

The CKG’s analytics core has an open modular design fully implemented in Python, exploiting open-source libraries that are widely employed, well maintained and that cover a broad data science ecosystem: statistics, network analysis, machine learning and visualization. Using these libraries ensures the quality, robustness and efficiency of the underlying algorithms and methods. As we have shown here, it also enables readily incorporation of new developments in data science, which can quickly be adapted to specifically support proteomics data analysis. Increasing concerns about reproducibility of scientific results (Baker and Penny, 2016; 2016) are also addressed by the CKG. All its algorithms are open source, databases are version controlled and there is full documention of the default and custom analyses. We employ Jupyter notebooks to generate shareable analysis pipelines that make results reproducible and replicable. Our diverse case studies show how these notebooks enable seamless multidimensional data integration and advanced analyses. We envision widespread adoption of this framework by the proteomic community and beyond.

Furthermore, the CKG has the potential to work as a clinical decision support tool and we have already used it in this capacity (Doll et al., 2018, 2019b). A next step could be the integration of electronic patient records (EHRs) into the graph, which has been suggested as a way to bridge the gap between research and direct patient care (Jensen et al., 2014; Nelson et al., 2019). However, this will require addressing a plethora of regulatory and ethical issues (Abul-Husn and Kenny, 2019; Grote and Berens, 2020). Medical institutions in different geographical areas often have different database systems and data formats, making data harmonization difficult. This challenge has been internationally recognized and standards such as the Fast Healthcare Interoperability Resources (FHIR) define how information contained in EHRs should be formatted (Boussadi and Zapletal, 2017). CKG can easily incorporate FHIR into its data model as the next step.

The different components of the CKG allow individual research groups to analyze, integrate and build a database of their proteomics and other omics projects. Reports and notebooks can readily be shared to replicate the analyses, thereby contributing to reproducible science (Figure 7A). Beyond this, the open nature and free availability of CKG also lend themselves to aggregate data and knowledge in what we term a community graph (Figure 7B). This would ensure that the community benefits from similar proteomics or omics projects performed elsewhere. For biomarker discovery this constitute an extension of the ‘rectangular strategy’ (Geyer et al., 2017), allowing direct and deep project comparison and leading to increasingly more robust and powerful analysis and knowledge generation.

**Figure 7.**
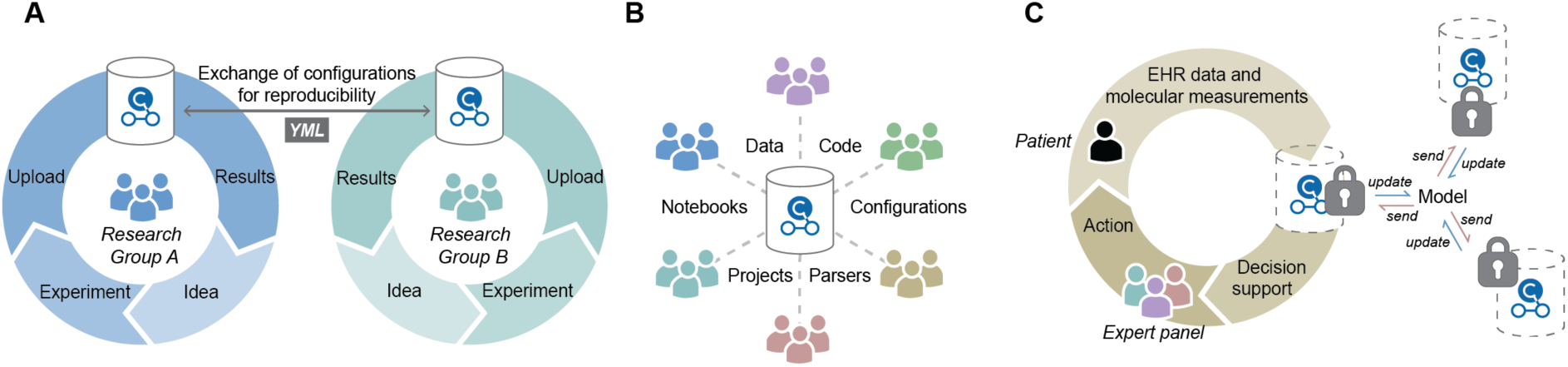
Future perspectives. A) Reports and notebooks in local graphs can readily be shared to replicate analyses, thereby contributing to reproducible science. B) Aggregating data and knowledge of multiple projects from different groups within a community would allow direct and deep project comparison and lead to increasingly more robust and powerful analysis and knowledge generation. C) To protect the sensitive nature of healthcare data and still allow researchers to train models and learn from the data the CKG could be implemented as a protected graph using federated learning.

We envision that different groups and institutions will have their own local version of the CKG, protecting the sensitive nature of healthcare data, but in a way that still enables cross-platform analyses. New approaches, such as differential privacy and federated learning (Bonawitz et al., 2019; Brisimi et al., 2018), would allow researchers to use the CKG to train models iteratively across institutions without direct access to the sensitive data (Figure 7C). The CKG could also integrate with existing standardized health data warehouse solutions, such as i2b2 and OMOP (Boussadi and Zapletal, 2017).

In conclusion, we describe the CKG, an open, roubust framework for transparent, automated and integrated analysis of proteomics and multi-level omics data, designed to incorporate all the prerequisites for reproducible science. Building on previous work of the Python and R scientific communities, it is itself meant to enable a community effort in which omics data and projects do not simply accrue in silos but actively contribute to a web of connected knowledge. Apart from the basic biomedical science context, it is especially designed for future application in the context of biomedicine as it provides a framework for harmonizing and integrating a diverse range of clinically relevant data and knowledge. The CKG thus directly addresses some of the major bottlenecks towards personalized medicine and rigorous, data-driven clinical decision-making process. We envision that based on this initial report, others in biomedical and clinical research community will be encouraged to contribute and further develop this platform into an easy to use tool implementable to directly support clinical decision-making.

## Methods

### Graph database

Graph databases are NoSQL databases that represent and store data using graph structures. The graph structure is a collection of nodes and edges, which represent relationships between the nodes, and properties. Storage of data in such structure facilitates access to densely connected data by providing graph-traversal linear times. The Clinical Knowledge Graph implements a graph database that contains more than 16 million nodes (33 labels) and more than 110 million relationships (45 different types). The database is built using Neo4j Community Edition (https://neo4j.com/), a scalable native graph database that allows storage, management and analysis of interconnected data. Neo4j provides a query language specific for graph structures, Cypher, and an extensive library of procedures and functions (APOC library and Graph Algorithms) that can be used for data integration, data conversion or graph analysis. Further, Neo4j makes the database available via several protocols (bolt, http or https) and provides a mission control center that interfaces with the database and helps manage it.

### Data integration

#### Ontologies

To build the Clinical Knowledge Graph database, we selected the different node labels (33 labels) and relationship types (45 types) between them to design the graph data model (Figure 3A). These nodes and relationships were defined based on the type of biological or clinical questions or problems set out to respond or solve. For each node label we defined the identifiers by using commonly used biomedical ontologies or standard terminologies. Ontologies denote concepts, in this case nodes (e.g. diseases) and provide an acyclic graph structure that describes how these concepts are related. We benefited from this underlying structure to integrate these concepts and relationships (‘is_a’ relationships) directly into the knowledge graph. Likewise, we integrated the terms and relationships standardized in terminologies such as SNOMED-CT, which defines clinical terms and their associative relationships.

Some of the nodes in our graph data model could not be described using ontologies or existing terminologies and they needed to be standardized using identifiers from the selected biomedical databases (e.g. UniProt, HMDB, etc) (Suppl Table S1).

#### Databases

Once the graph data model and the node label identifiers were defined, we selected multiple well-known and used biomedical databases (25 databases) (Suppl Table S1) to feed the Clinical Knowledge Graph. The selection of databases to be integrated responded to the type of nodes and relationships in the model and was also based on criteria such as access, usability, stability and acceptance by the research community. However, the flexible design of the graph database and the Clinical Knowledge Graph platform allows quick integration of new databases, ontologies, terminologies or even modifications in the data model (new nodes or relationships) (see methods GraphDB builder).

#### Experiments

The Clinical Knowledge Graph database models multiple node types, which in principle allows integration of different data types: genomics, transcriptomics, proteomics or metabolomics. However, the focus of the graph is initially on integration of quantitative mass spectrometry-based proteomics data, which may have influenced the structure of the data model specifically on how experimental projects are defined and stored. Similarly, the clinical context in which the database has been built limits the data to Human, while other species are not covered by the graph yet.

Proteomics data can be integrated by creating a new project, which requires defining new nodes in the database: enrolled experimental subjects, biological samples collected from these subjects and analytical samples extracted from those biological samples. Analytical samples correspond to the actual sample analysed in the Mass Spectrometer. All these nodes will have external identifiers and they will be mapped to unique internal identifiers in the knowledge graph. Internal identifiers will then be used to integrate experimental and clinical data seamlessly.

The relationship between analysed samples and proteins ((Analytical_sample)-[:HAS_QUANTIFIED]-(protein)) where the quantification (LFQ intensity) is stored as a property/attribute of the relationship (value) is obtained from proteomics data processed in advance. Currently, MaxQuant and Spectronaut output files or tabular files can be automatically loaded into the database using specific configuration (YAML file) for each format.

Similarly, clinical data – clinical variables collected for each subject or biological sample (in case of longitudinal studies) – can also be automatically loaded into the database in tabular format. As a requirement, all clinical variables provided need to follow the SNOMED-CT standard.

### Clinical Knowledge Graph Platform

#### Software Architecture

The Clinical Knowledge Graph platform has been designed using a modular architecture that divides the platform into functional compartments: graphdb_connector, graphdb_builder, report_manager and analytics_core (Figure 1A). Each module can be used independently, which provides a flexible environment to cover different scenarios and different needs: direct programmatic interaction with the database, deployment of local knowledge graph database, visualization of automatically analysed data from the database or just data analysis and visualization.

In combination, all modules provide a full workflow from project ideation and creation to analysis and visualization of results (Suppl. Figure S1). Additionally, we have included Jupyter notebooks as another layer of functionality, which allow further and specific analyses and serve as a playground for continuous improvement of the analysis and visualization functionality. Furthermore, notebooks will support replicability, reproducibility and reusability of analysis in the Clinical Knowledge Graph.

All the modules have been developed in Python 3.6. Some of the analyses are performed using R packages (e.g. SAMR and WGCNA) called from Python using the Rpy2 library. The library version used in the Clinical Knowledge Graph (rpy2== 3.0.5) is not compatible with Windows and these analyses are not available in installations in this operating system. Alternatively, we have created a Dockerfile, which holds all the necessary instructions to generate a complete container with all the requirements, in this setup windows users have all analyses available. When running the Docker container, four ports will be available 1) Neo4j HTTP port (7474), 2) Neo4j bolt port (7687) 3) CKG Dash server (8050) and 4) JupyterHub server (8090). The entry point to the container (docker_entrypoint.sh) defines all the steps needed: clone the gitHub repository, start the required services (Neo4j, JupyterHub, redis, celery), generate the database using the graphdb builder and running the report manager dash app.

All the code can be accessed at https://github.com/MannLabs/CKG. For further instructions on each functional module refer to the documentation at https://CKG.readthedocs.io.

#### GraphDB Connector

The graphdb connector is a functional module that provides functionality to connect and query the Clinical Knowledge Graph database. This module is Neo4j-dependent, it uses the Python library py2neo, but independent from the other functionality in the platform, which allows an agnostic interaction with the database and facilitates adaptation and scalability. Likewise, queries to the database in Cypher language across the platform have been defined as YAML objects with a structure that makes them findable (name, involved nodes and relationships), understandable (description) and easily replaceable.

#### GraphDB Builder

This functional module can be used to generate the Clinical Knowledge Graph database. It is divided into two steps: importing and loading. The import (importer.py) downloads the ontologies, terminologies and biomedical databases into the data directory, and formats the data into tabular files (nodes and relationships). The tabular files created by the importer are also stored in the data directory under the imports folder and organized into: ontologies, databases and experiments. Further, the import step generates some statistics (Hierarchical Data Format, HDF) regarding the number of nodes and relationships formatted as well as file sizes for each ontology, database or experiment. These statistics can be used to track possible errors in the import (data/imports/stats).

Once the import process finishes, data can be loaded into the graph database by the loader (loader.py), which runs several Cypher queries defined as YAML objects (cypher.yml) and loads the tabular files located in the import folder into the running database. To facilitate this two-step process, we have implemented a module called builder (builder.py), which can be used to either perform both steps or, one or the other. This module also allows importing or loading of specific ontologies, databases or experiments. After running the two steps, the running database should contain all the nodes and relationships harmonized from the different sources of data.

#### Analytics Core

Analytics core is divided into two main functionalities: analytics and visualization. Both modules are independent of the Clinical Knowledge Graph database and can be used to analyze and/or visualize data in the shape of a dataframe (Pandas dataframe). The analytics functionality uses Python statistics and data science libraries to implement the state-of-the-art analyses of proteomics data – data processing, differential expression, and correlation, enrichment and network analyses – and incorporates some recent relevant methods such as Weighted Gene Co-expression Network Analysis (WGCNA) or Similarity Network Fusion analysis (SNF). Moreover, to ensure the correct use of these functions, they are designed to identify the experimental design automatically and consequently define the appropriate statistical analysis to perform (e.g. dependent or independent test). The visualization library (viz) uses Plot.ly, an interactive graphing library for Python and R, which opens the possibility to save plots in a format compatible with both programming languages (JSON format).

#### Report Manager

The Report manager is a tool to interface with the existing projects in the Clinical Knowledge Graph database. This functional module makes use of the Analytics core to analyze the project data and to generate interactive graphs, and then create detailed reports with these analyses. These reports can be accessed through dashboard apps implemented in Plot.ly Dash (https://plot.ly/dash/). The dash server can be started by running the index module (index.py), and accessed at http://localhost:5000. The initial app (home) redirects to the login page if there are users in the database, otherwise it shows the current data model and statistics about the database such as number of nodes and relationships of each type. Further, this app also links to the other existing pages: project creation, data upload and imports, and lists all the existing projects in the database. When a link to an existing project is accessed for the first time the report manager runs the automated analyses for each data type in the project using the default configuration. Reports for each data type are shown in tabs in the Project app and two extra tabs are also present: the multiomics tab, if there is more than one data type (e.g. clinical and proteomics data), and the knowledge graph tab, which shows a summary figure of all the other tabs.

New report analysis pipelines can be defined using configuration files (YAML format) describing the arguments to be used in the data processing, as well as the sequence of analyses to be performed. The structure requires, for each analysis’ configuration, to specify which data to use (name of the dataframe/s), a list of analyses and plots to visualize results (functions in the analytics core: analytics and viz respectively), whether or not to store the results as dataframes, and the arguments needed for analysis and visualization.

Once generated, project reports are stored in HDF5 format so that they can be quickly shown when accessed again. However, project reports can be regenerated either with default configuration or by providing specific configuration files using the ‘Change Analysis’ Configuration’ option in the Project app. The saved reports can also be easily accessed programmatically with functionality within the report manager (project.load_project_report()) or using the R library rhdf5.

Reports can be downloaded in a compressed file (zip), which contains one folder for each generated tab and inside, all the dataframes created during the analyses (tab separated format, tsv), all the plots as vector graphics and all networks in Graph Modeling Language (gml) compatible with Cytoscape and JSON (Christmas et al., 2005).

#### Notebooks

We included Jupyter notebooks as another component of the Clinical Knowledge Graph platform. This component serves a two-fold purpose 1) playground to test and develop new analyses and visualizations and 2) as a collection of case studies that can be shared, reproduced, replicated and reused.

We have defined several Jupyter notebooks that show functionality and analyses that broaden the scope of the report manager. These notebooks show: how to work with reports in R, how to extend the default analysis in the report, and how to analyze external data making use of the Clinical Knowledge Graph platform (Archer et al., 2018).

## QUANTIFICATION AND STATISTICAL ANALYSIS

### Case studies

#### Non-alcoholic Fatty Liver Disease Study

We use a previously published internal proteomics dataset (Niu et al., 2019) (PXD011839) as a showcase of the capabilities of the Clinical Knowledge Graph. In this publication, Niu et al studied the plasma proteome profiles of 48 patients with and without cirrhosis or Non-Alcoholic Fatty Liver Disease (NAFLD) and identified several statistically significantly changing proteins, some of which were already linked to liver disease. We aimed to reproduce the results obtained using the automated default analysis pipeline of the Clinical Knowledge Graph.

#### Downstream Rapid Proteomics Analysis

We use a previously published internal proteomics dataset (Doll et al., 2018) (PXD008713). This study presents a rapid proteomics analysis that identified a possible alternative treatment for an end-stage cancer patient. We built a downstream analysis pipeline to accelerate and prioritize alternative candidate drug treatments using the Clinical Knowledge Graph. We provide a Jupyter Notebook to show how functionality implemented in the graphdb_connector module (query_utils.py) can be used to single out queries that can help find known links between identified upregulated proteins and inhibitory drugs, between those drugs and known side effects and publications, and how to use this knowledge to prioritize drug candidates.

#### Multi-level Proteomics Analysis

We reanalyzed and extended a multi-level proteomics study, including interactomics and phosphoproteomics, that provides insights into the mechanisms of resistance to platinum-based chemotherapy in high-grade ovarian serus adenocarcinoma (Coscia et al., 2018) (PXD010372). The CKG reproduce the findings and extend them with deeper analysis of the protein complexes identified (Giurgiu et al., 2019) and substrate and phosphosite-specific annotations (Hornbeck et al., 2015; Perfetto et al., 2016).

### Clinical Knowledge Graph Update

Databases and ontologies integrated in the CKG can be updated in two different modes using the graphdb_builder. The builder can run a full update, which will regenerate the entire database with newly downloaded data from the sources, or a partial update by specifying what sources to be imported and loaded into the graph. The partial update can also be used to extend the graph when a new database or ontology is added.

Experiments can be updated using the ’Data Upload’ functionality in the dashboard app by indicating the project identifier and uploading the new data. When a full update is performed in CKG’s graph which involves upgrading the version of essential databases such as UniProt (UniProt Consortium, 2018), it is highly recommended to process the RAW proteomics data searching with the new version of the proteome and to generate again all the project reports with the new data.

CKG’s is an open source project and its code will continue to grow and improve through version control in the GitHub repository (https://github.com/MannLabs/CKG). Currently, version 1.0.0 is available and new releases will be made available in a controlled manner and named following the PEP 440 specification (https://www.python.org/dev/peps/pep-0440/). CKG is an open source project and contributions will help the framework to grow with additional ontology, database or experimental parsers, improved documentation, increased testing and feedback. Specific details on how to contribute can be found in CKG’s documentation.

### Installation and Hardware Requirements

CKG’s purpose and architecture define it as a multi-user platform that requires installation in a server-like setup and administration knowledge. However, individual users can have local installations making sure hardware and software requirements are fullfiled. For specific requirements, consult CKG’s documentation at https://CKG.readthedocs.io.

## DATA AND CODE AVAILABILITY

All code is available at https://github.com/MannLabs/CKG. and detailed documentation can be found at https://CKG.readthedocs.io.

## Acknowledgments

We thank all members of the Proteomics and Signal Transduction Group (Max Planck Institute) and the Clinical Proteomics Group (Novo Nordisk Foundation Center for Protein Research), especially Eugenia Voytik, Fabian Coscia, Lisa Schweizer, Lili niu and Niels Skotte for their help testing the code and providing feedback. This project was supported by Novo Nordisk Foundation grants (NNF14CC0001 and NNF15CC0001). F.C acknolwedges the European Union’s Horizon 2020 research and innovation program (Marie Skłodowska-Curie individual fellowship under grant agreement 846795). We thank the Python community for the excellent scientific libraries developed and maintained and also Neo4j for providing a community version of their graph database and its community for helping improve the platform.

## Author Contributions

A.S., R.C., and A.B.N. implemented CKG’s code. A.S., M.M. conceived and designed the framework. A.S. R.C., L.N., P.E.G., F.C. and A.B.N contributed to the data analysis for the case studies presented. L.J.J, NJWA, P.E.G. and F.M. contributed with data and expertise on how to integrate them. A.S., R.C., A.B.N. and M.M. wrote and edited the manuscript and the figures. A.S. and M.M. directed the study.

## Supplemental Information

Supplementary tables (S1,S3, S4) - Suppl tables.xlsx

Supplementary table S2 – TableS2.pdf

Supplementary Figures - Suppl Figures.pdf

