## Supplementary Figures S1-3 for "Clinical Knowledge Graph Integrates Proteomics Data into Clinical Decision-Making"

**Figure S1**

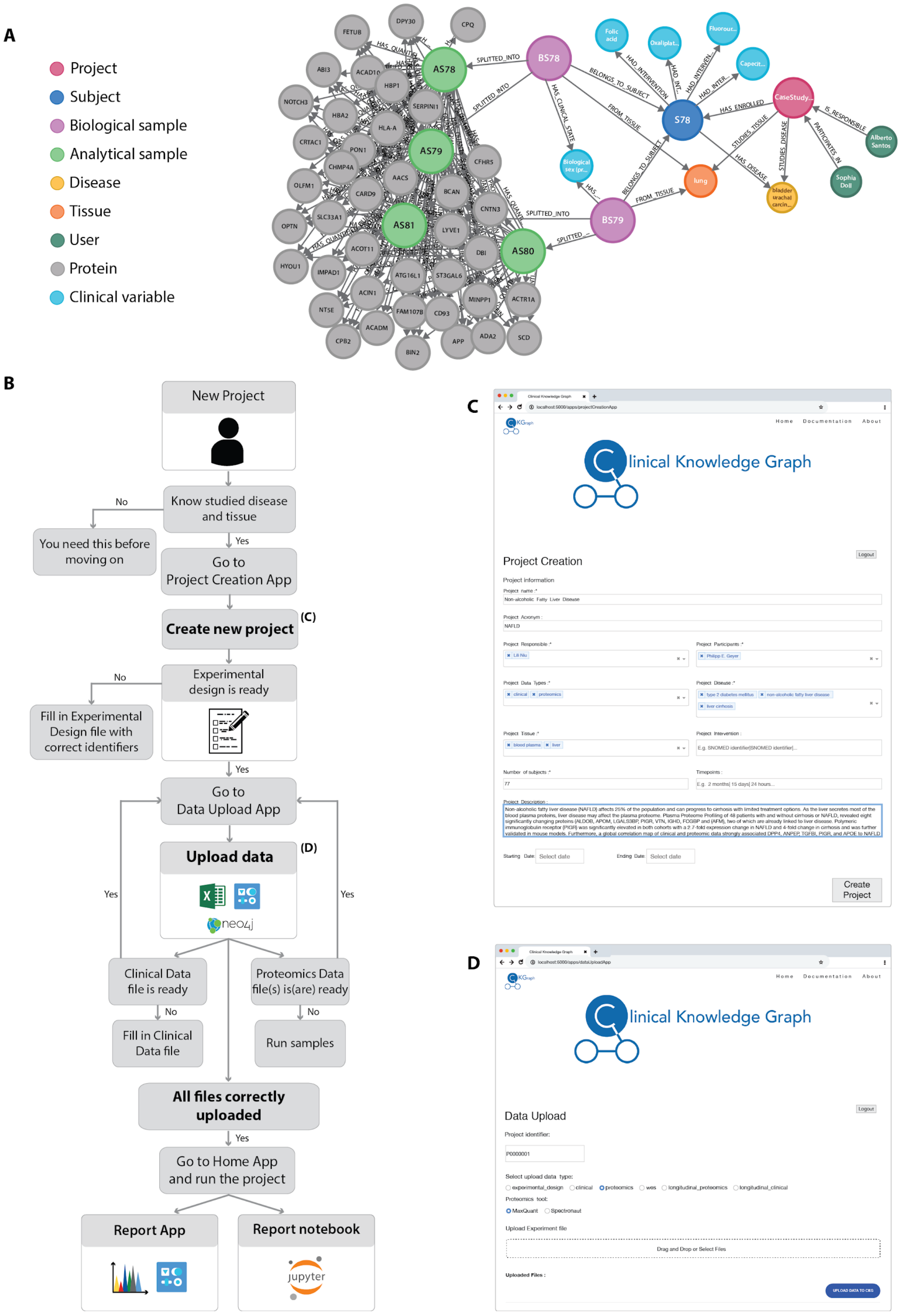

**Figure S1.** A) CKG Neo4j graph structure showing how metadata is stored around a research project. Nodes for project, subject, biological sample, analytical sample and quantified proteins are depicted, together with the relationships between them. B) Workflow from project idea to knowledge-based analysis report. Once a project is created, relevant data files can be uploaded, starting with the experimental design file, and followed by the clinical data and proteomics files. When the upload is finished successfully, the user can navigate to the homepage app and run the default analysis pipeline by selecting the respective project. C) Example of how the Project Creation App looks like, and how it should be filled in. D). Data Upload App. Type in the correct project id, select the data type to be uploaded and the appropriate files. Once all files have been selected, press the bottom button to upload the data.

### Figure S2

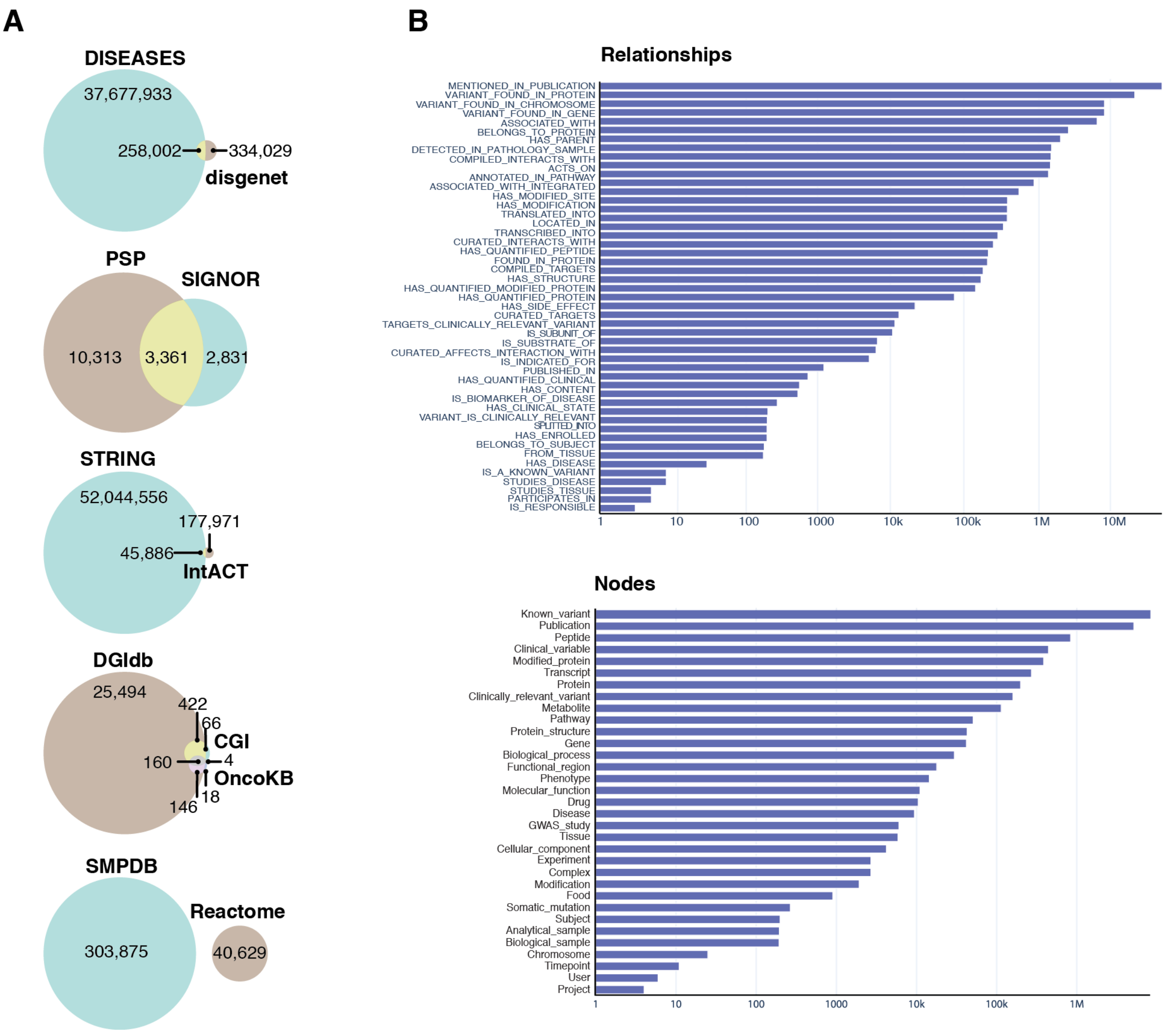


**Figure S2.** Distributions of nodes, relationships and overlapping sources.
A) Venn diagrams showing the number of relationships originating from different sources, for the five cases where information from multiple databases are used to obtain one relationship type.
B) Barplots of the number of times each relationship- and node type appear in the graph database, respectively.

### Figure S3

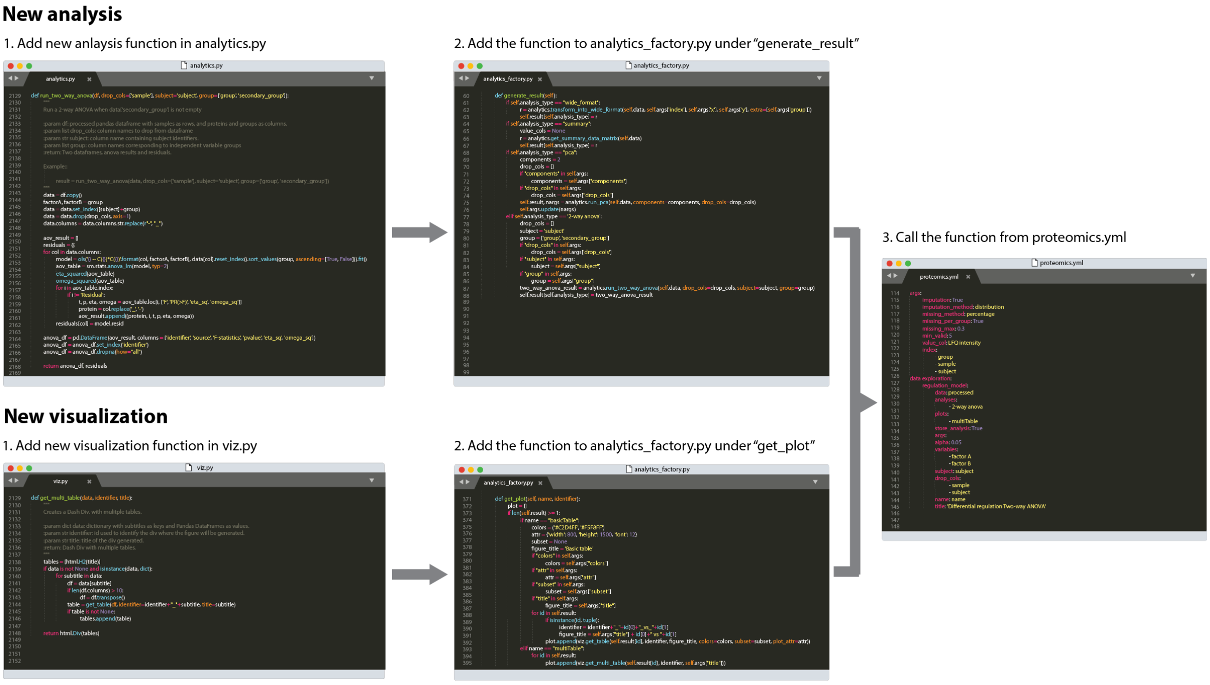


**Figure S3.** Adding new functions to the CKG. New analysis and/or visualizations can be added to the CKG pipeline in three simple steps: adding the new function to the appropriate *analytics_core* module, adding a call for the function in the *analytics_factory* module and finally, calling it in the appropriate configuration file.

#### Table S1. Databases

List of databases integrated into the Clinical Knowledge Graph database.

#### Table S2. Analytics core

All the analysis and visualization functions available in the analytics core.

#### Table S3. Link Prediction

Predicted mapping between Gene Ontology biological processes and Reactome metabolic pathways based on similar protein and metabolite annotations.

#### Table S4. Protein-drug relationships

Number of protein-drug relationships and the effect of the drugs on these proteins according to STITCH database (http://stitch.embl.de/).
