## Supplementary material for "Clinical Knowledge Graph Integrates Proteomics Data into Clinical Decision-Making": Table S2

| | Step | Method | Description | CKG function | Reference | Link |
|---|---|---|---|---|---|---|
| Data preparation | Filtering | percentage | Filtering based on maximum percentage of missing values allowed (per group optional) | extract_percentage_missing | | |
| | | at_least_x | Filtering based on minimum number of present values (per group optional) | extract_number_missing | | |
| | Imputation | K-nearest Neighbors | Imputation based on the algorithm Nearest Neighbors (NN) | imputation_KNN | https://www.ncbi.nlm.nih.gov/pubmed/21743766 | https://pypi.org/project/fancyimpute/ |
| | | Probabilistic Minimum Imputation approach | Imputation method replacing missing values with values withdrawn from a down-shifted normal distribution | imputation_normal_distribution | https://www.ncbi.nlm.nih.gov/pubmed/26906401 | |
| | | Mixed model | A combination of KNN and Probabilistic Minimum depending on the number of values available (>60% KNN, rest ProbMin) | imputation_mixed_norm_KNN | | |
| | Normalization | Median | Normalize samples using the median | median_normalization | | |
| | | Median polish | Normalization based on the medians obtained from the rows and the columns to iteratively calculate the row effect and column effect on the data | median_polish_normalization | | |
| | | Quantile | Adjustment method that forces the observed distributions to be the same and it uses the average of each quantile across samples as the reference (assumes the statistical distribution of each sample is the same) | quantile_normalization | | |
| | | Linear | Apply l1 or l2 normalization | linear_normalization | | https://scikit-learn.org/stable/modules/preprocessing.html |
| Data exploration | Ranking | Ranking | Ranking of proteins based on intensity | get_ranking_with_markers | | |
| | Coefficient of Variation | Coefficient of Variation | Coefficient of variation per group as a quality control | get_coefficient_variation | | |
| | QC markers | QC Makers | If there are quality control markers associated with the tissue studied, they are used to visualize possible outliers (z-score) among the samples | run_qc_markers_analysis | https://www.ncbi.nlm.nih.gov/pubmed/31566909 | |
| | Summary statistics | Summary statistics | Statistics for rows and columns in the data matrix | get_summary_data_matrix | | |
| Data analysis | Dimensionality reduction | PCA | Principal Component Analysis (2D, 3D) | run_pca | https://www.ncbi.nlm.nih.gov/pubmed/24061923 | https://scikit-learn.org/stable/modules/generated/sklearn.decomposition.PCA.html |
| | | tSNE | t-distributed Stochastic Neighbor Embedding | run_tsne | https://www.ncbi.nlm.nih.gov/pubmed/30252473 | https://scikit-learn.org/stable/modules/generated/sklearn.manifold.TSNE.html |
| | | UMAP | Uniform Manifold Approximation and Projection | run_umap | https://arxiv.org/abs/1802.03426 | https://umap-learn.readthedocs.io/ |
| | Hypothesis testing | SAMR | Significance analysis of microarrays applied to proteomics data | run_samr | https://www.ncbi.nlm.nih.gov/pubmed/11309499 | https://www.rdocumentation.org/packages/samr/versions/3.0 |
| | | ANOVA | Analysis of Variance | run_anova | | https://pingouin-stats.org/generated/pingouin.anova.html |
| | | ANOVA-rm | Analysis of Variance for repeated measurements | run_repeated_measurements_anova | | https://pingouin-stats.org/generated/pingouin.rm_anova.html |
| | | t-test | t-test mean difference | run_ttest | | https://pingouin-stats.org/generated/pingouin.ttest.html |
| | Multiple-test correction | Bonferroni | Bonferroni p-value correction | apply_pvalue_correction | | https://www.statsmodels.org/stable/generated/statsmodels.stats.multitest.multipletests.html |
| | | Benjamini-Hochberg | Benjamini-Hochberg FDR correction | apply_pvalue_correction | | https://www.statsmodels.org/stable/generated/statsmodels.stats.multitest.multipletests.html |
| | | Permutation-FDR | Permutation FDR | apply_pvalue_permutation_fdrcorrection | https://www.jstor.org/stable/2346101 | |
| | | ... | ... | apply_pvalue_correction | | https://www.statsmodels.org/stable/generated/statsmodels.stats.multitest.multipletests.html |
| | Correlation | Correlation | Pearson or Spearman correlation | run_correlation | | |
| | | Correlation-rm | Pearson or Spearman correlation for repeated measurements | run_rm_correlation | | https://pingouin-stats.org/generated/pingouin.rm_corr.html |
| | Enrichment | Fisher Exact Test | Significant test for contingency tables | run_enrichment | | https://docs.scipy.org/doc/scipy/reference/generated/scipy.stats.fisher_exact.html |
| | Network analysis | Louvain partition | Best partition algorithm | get_louvain_partitions | https://www.ncbi.nlm.nih.gov/pubmed/21517554 | https://python-louvain.readthedocs.io/en/latest/api.html |
| | | Greedy modularity | A hierarchical agglomeration algorithm for detecting community structure | get_network_communities | https://www.ncbi.nlm.nih.gov/pubmed/15697438 | https://networkx.github.io/documentation/stable/reference/algorithms/generated/networkx.algorithms.community.modularity_max.greedy_modularity_communities.html |
| | | Asynchronous label propagation algorithm | The algorithm initializes each node with a unique label and repeatedly sets the label of a node to be the label that appears most frequently among that nodes neighbors. The algorithm halts when each node has the label that appears most frequently among its neighbors | get_network_communities | https://www.ncbi.nlm.nih.gov/pubmed/17930305 | https://networkx.github.io/documentation/stable/reference/algorithms/generated/networkx.algorithms.community.label_propagation.asyn_lpa_communities.html#id3 |
| | | Girvan-Newman algorithm | Hierachical algorithm that removes edges iteratively and defining communities by the remaining connected components | get_network_communities | | https://networkx.github.io/documentation/stable/reference/algorithms/generated/networkx.algorithms.community.centrality.girvan_newman.html |
| | | Affinity propagation | A centroid-based clustering algorithm that finds members of the input set that are representative of clusters and estimates the number of clusters | get_network_communities | https://www.ncbi.nlm.nih.gov/pubmed/17218491 | https://scikit-learn.org/stable/modules/generated/sklearn.cluster.affinity_propagation.html |
| | Multiomics | WGCNA | Weighted gene co-expression network analysis for describing the correlation patterns among proteinsfinding clusters (modules) of highly correlated genes, for summarizing such clusters using the module eigengene or an intramodular hub gene, for relating modules to one another and to external sample traits | run_WGCNA | https://www.ncbi.nlm.nih.gov/pubmed/19114008 | https://horvath.genetics.ucla.edu/html/CoexpressionNetwork/Rpackages/WGCNA/ |
| Visualization | Viz | Pie chart | Circular statistical chart, which is divided into sectors to illustrate numerical proportion | get_pieplot | | https://plotly.com/python/pie-charts/ |
| | | Distribution plot | Representations of statistical distributions | get_distplot | | https://plotly.com/python/distplot/ |
| | | Bar chart | | get_barplot | | https://plotly.com/python/bar-charts/ |
| | | Scatter plot matrix | | get_facet_grid_plot | | |
| | | Ranking plot | | get_ranking_plot | | |
| | | Scatter plot | | get_simple_scatterplot | | |
| | | Volcano plot | | run_volcano | | |
| | | Heatmap plot | | get_heatmapplot | | |
| | | Heatmap plot with annotation and clustering | | get_complex_heatmapplot | | |
| | | Network | Generates a Cytoscape network (Plot.ly), a Jupyter notebook compatible Cytoscape network (Cyjupyter) and a json format network | get_network | https://www.ncbi.nlm.nih.gov/pubmed/14597658 | https://dash.plotly.com/cytoscape |
| | | PCA plot | PCA plot with loadings (2D and 3D) | get_pca_plot | | |
| | | Sankey diagram | Visualize the contributions to a flow | get_sankey_plot | | |
| | | Table | | get_table | | https://dash.plotly.com/datatable |
| | | Violin plot | | get_violinplot | | https://plotly.com/python/violin/ |
| | | Parallel coordinates plot | | get_parallel_plot | | https://plotly.com/python/parallel-coordinates-plot/ |
| | | WGCNA plots | Generates all the plots for the WGCNA analysis | get_WGCNAPlots | https://www.ncbi.nlm.nih.gov/pubmed/19114008 | https://horvath.genetics.ucla.edu/html/CoexpressionNetwork/Rpackages/WGCNA/ |
| | | 2-way Venn diagram | | get_2_venn_diagram | | https://plotly.com/python/shapes/ |
| | | Word cloud | Represents the frequency of words in a text using size and color | get_wordcloud | | https://github.com/PrashantSaikia/Wordcloud-in-Plotly |
| | | Save Dash plot | Save a Dash figure object to svg format | save_DASH_plot | | |
| | | Kaplan-meier plot | Kaplan-meier survival plot with significance annotation | get_km_plot | | https://plotly.com/python/v3/ipython-notebooks/survival-analysis-r-vs-python/ |
| | | Polar chart | Represents data along radial and angular axes | get_polar_plot | | https://plotly.com/python/polar-chart/ |
